# Drug Metabolism and Transport Capacity of Endothelial Cells, Pericytes, and Astrocytes: Implications for CNS Drug Disposition

**DOI:** 10.1101/2024.08.01.606165

**Authors:** Hannah N. Wilkins, Stephen A. Knerler, Ahmed Warshanna, Rodnie Colón Ortiz, Kate Haas, Benjamin C. Orsburn, Dionna W. Williams

## Abstract

Therapeutically targeting the brain requires interactions with endothelial cells, pericytes, and astrocytes at the blood brain barrier (BBB). We evaluated regional and cell-type specific drug metabolism and transport mechanisms using rhesus macaques and *in vitro* treatment of primary human cells. Here, we report heterogenous distribution of representative drugs, tenofovir (TFV), emtricitabine (FTC), and their active metabolites, which cerebrospinal fluid measures could not reflect. We found that all BBB cell types possessed functional drug metabolizing enzymes and transporters that promoted TFV and FTC uptake and pharmacologic activation. Pericytes and astrocytes emerged as pharmacologically dynamic cells that rivaled hepatocytes and were uniquely susceptible to modulation by disease and treatment. Together, our findings demonstrate the importance of considering the BBB as a unique pharmacologic entity, rather than viewing it as an extension of the liver, as each cell type possesses distinct drug metabolism and transport capacities that contribute to differential brain drug disposition.

## Introduction

The global impact of neurologic disease is astounding as it affects every developmental stage and virtually all facets of life, including movement, cognition, and the capacity for human connection. For these reasons, substantial drug discovery efforts focus on developing small molecule therapeutics whose targets have implications in prevention and treatment of neurologic disease. Yet, even when putative therapies with great clinical promise are identified, the blood brain barrier (BBB) creates a major obstacle for central nervous system (CNS) penetration that poses significant challenges in achieving pharmaceutical efficacy ^1^. As such, understanding the mechanisms underlying drug distribution in the brain is poised to have a wide-reaching impact on the millions of lives impacted by neurologic disease.

While endothelial cells encompass the primary portion of the brain microvasculature, the essentiality of the surrounding neurovascular unit in maintaining appropriate BBB function cannot be ignored. Emerging evidence demonstrates that pericytes and astrocytes are more than support cells and play distinct roles from endothelial cells, without which appropriate BBB function would not occur ^2,3^. Even still, understanding of pericyte and astrocytic contributions to the BBB are primarily restricted to permeability and integrity. As such, their capacity to contribute to CNS drug disposition is not widely considered. This limitation becomes even more apparent due to the regional variability that exists at the BBB, wherein diverse anatomic sites have differential pericyte and astrocyte innervation that is dynamically regulated ^4^. The relative contribution of endothelial cells, pericytes, and astrocytes to regional heterogeneity in CNS drug distribution remains unknown, the understanding of which will be important to identify mechanisms to target drug to specific anatomic sites in the brain.

We aimed to address this question by evaluating drug metabolizing enzymes and membrane-associated transporters as they are essential for drug efficacy, biotransformation, elimination, and distribution in tissues. These pharmacological pathways of drug metabolism and transport are primarily characterized in peripheral organs, including liver, yet remain largely unknown at the BBB. Our study used antiretroviral therapies (ART) primarily used for the prevention and treatment of human immunodeficiency virus-1 (HIV) to probe this question as they are uniquely positioned to serve as an investigative model for CNS drug disposition. The CNS represents one of the most challenging viral sanctuaries for HIV as it invades the brain within two weeks of infection resulting in significant neurologic consequences in ∼50% of people living with the virus ^5,6^.

Here, we used the first-line nucleot(s)ide reverse transcriptase inhibitors, tenofovir (TFV) and emtricitabine (FTC), to 1) evaluate regional differences in drug distribution across the brain and 2) elucidate differential drug transport and metabolizing capabilities of brain endothelial cells, pericytes, and astrocytes, which may serve as a model of CNS drug disposition pathways that determine drug access at the dynamic microenvironment of the BBB. With a combination of *in vivo* studies in simian immunodeficiency virus (SIV)-infected and ART-treated rhesus macaques and *in vitro* HIV exposure and ART treatments of cell monocultures, we evaluated the capacity of each cellular component of the BBB to uptake and convert TFV and FTC into their pharmacologically active metabolites using a combination of liquid chromatography/mass spectrometry (LC-MS/MS), proteomics by mass spectrometry, and drug transporter and metabolizing enzyme functional assays. Our findings demonstrate that all cells at the BBB possess drug metabolizing and transporting capabilities that can contribute to CNS drug bioavailability and efficacy, which may inform pharmacodynamics, drug targeting, and drug discovery. Further, our work indicates that the BBB is a dynamic, pharmacologically active microenvironment that responds differentially to disease and CNS small molecule therapeutics in a cell-dependent manner.

## Results

### TFV and FTC Are Heterogenously Distributed Among Brain Regions and BBB Cell Types

It is well accepted that ART penetrates the brain to a lesser extent than peripheral tissues. Previously, ART cerebrospinal fluid (CSF) measurements were used as surrogates for brain tissue concentrations ^7^. However, measurements within brain tissue demonstrated a lack of concordance with CSF concentrations ^8,9^. These studies indicate that first-line ART drugs, TFV and FTC, are CNS-penetrant; however, the mechanisms at the BBB that facilitate their entry into the brain and contribute to heterogenous CNS distribution remain poorly understood ^8–10^. To address this, we obtained brain from SIV-infected rhesus macaques that received a once daily, 6-month ART regimen that included TFV and FTC and LC-MS/MS. We focused on thalamus, frontal cortex, and cerebellum as HIV infection of these brain regions is well-characterized and the resultant neurologic deficits that occur represent an important public health consideration ^11,12^. We found that TFV concentrations were higher in all three tissues relative to FTC, which was below the limit of quantification for all brain regions, except the thalamus (**Figure 1A**). Interestingly, TFV was heterogeneously present in the brain wherein it was 78% higher in the cerebellum (0.0784 ± 0.0448 ng/mg) relative to the thalamus (0.0438 ± 0.0619 ng/mg) and 32% higher than the frontal cortex (0.0595 ± 0.0842 ng/mg).

**Figure 1.**
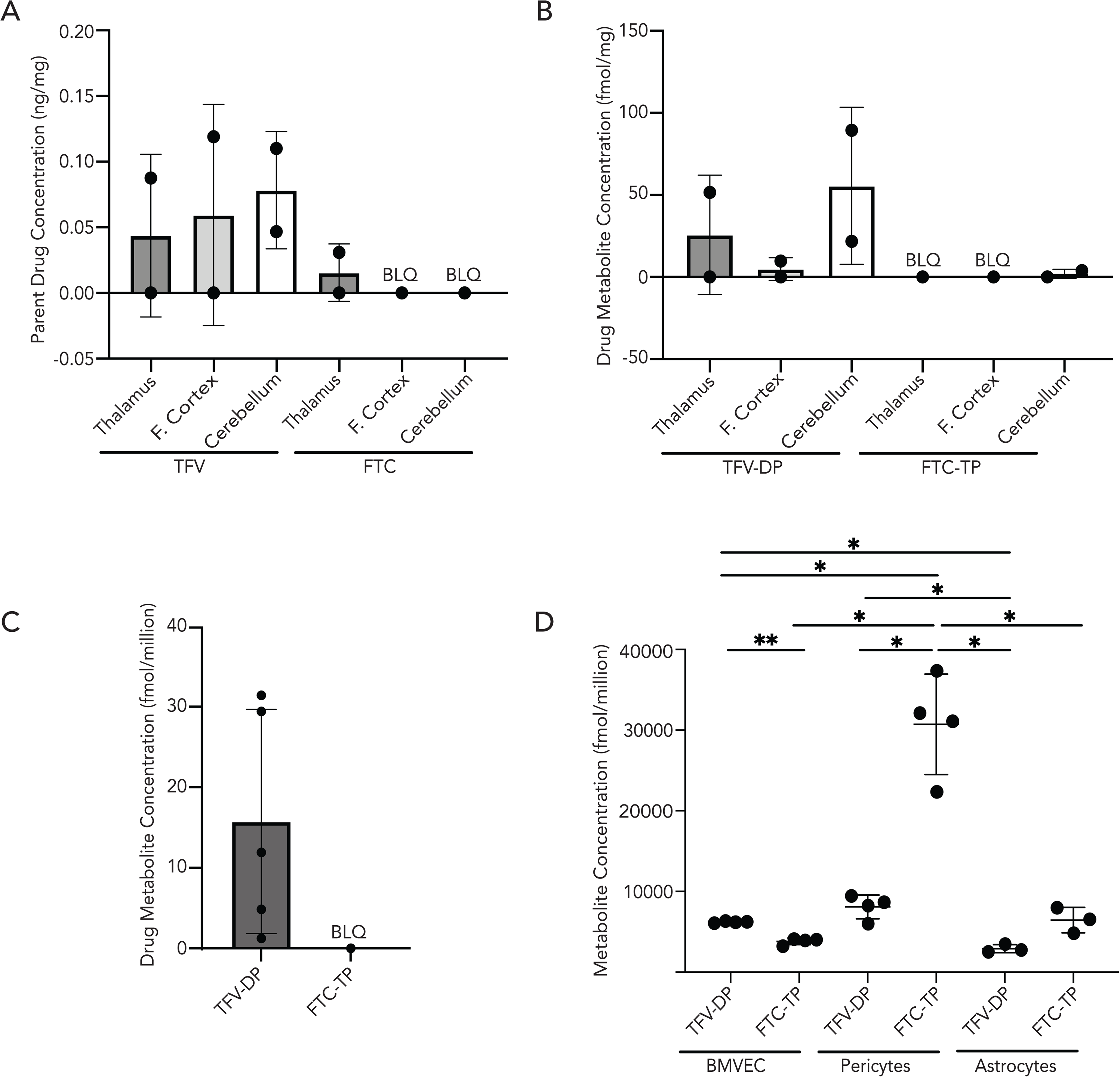
TFV and FTC parent and metabolite quantification in BBB cells and rhesus macaque tissues. **(A and B)** The thalamus, frontal cortex (denoted as F. Cortex), and cerebellum were sectioned from rhesus macaque brains collected 200 days post inoculation with SIVmac251 followed by a subcutaneous dose of 20 mg/kg TFV, 40 g/kg FTC, and 2.5 g/kg DTG administered once daily. Within these brain tissues, **(A)** Parent TFV and FTC as well as **(B)** TFV-DP and FTC-TP metabolites were quantified by LC-MS/MS from cerebellum, frontal cortex, and thalamus (n=2 macaques, represented as individual dots). **(C)** Astrocytes were obtained from healthy rhesus macaques that received six months of a once daily intramuscular ART administration (5.1 mg/kg TFV, 50 mg/kg FTC, and 2.5 mg/kg DTG). Astrocytes were isolated from five macaque brains (represented as individual dots) and TFV-DP and FTC-TP quantified by LC-MS/MS. Below limit of quantification (BLQ) samples reported (represented as a singular dot) had concentrations less than 5 fmol/sample (TFV-DP), 50 fmol/sample (FTC-TP), 0.05 ng/sample (TFV), and 0.25 ng/sample (FTC). **(D)** Primary human BBB cells were treated with 10 μM TFV, 2 μM FTC, and 6 μM DTG for 24h at 37°C, 5% CO_2_. After this time, TFV-DP and FTC-TP were quantified by LC-MS/MS. Three to four independent experiments were performed (represented as individual dots). Data represented as mean ± standard deviation. Statistical analysis was performed using Brown-Forsythe and Welch ANOVA. *p < 0.05, **p < 0.01, ****p < 0.0001.

We next evaluated the active metabolites of TFV and FTC, tenofovir-diphosphate (TFV-DP) and emtricitabine-triphosphate (FTC-TP), respectively, by LC-MS/MS, as these are the only versions of these drugs capable of suppressing HIV (**Figure 1B**).

TFV-DP concentrations were 116% and 1134% higher in the cerebellum (55.60 ± 47.80 fmol/mg) relative to the thalamus (25.75 ± 36.42 fmol/mg) and the frontal cortex (4.90 ± 6.92 fmol/mg), respectively, suggesting a differential capacity exists for pharmacologic activation across brain regions (**Figure 1B**). Further, TFV-DP (55.60 ± 47.80 fmol/mg) was 2866% higher than FTC-TP (1.94 ± 2.74 fmol/mg) in the cerebellum, highlighting a differential accumulation of ART active metabolites within brain regions. Similar to that which occurred for its parent drug, FTC-TP was below the limit of quantification for all brain regions, except the cerebellum. We did not detect TFV, FTC, or their active metabolites in brain obtained from untreated, control rhesus macaques (data not shown).

TFV and FTC entry into brain requires important interactions with the BBB. It remains unclear if, during these interactions, ART is retained within the cells comprising the BBB, or if instead they are transported in their entirety into the brain parenchyma. This is particularly important as astrocytes and pericytes are permissive to HIV and infection dysregulates their essential contributions to BBB function ^13,14^. While astrocyte uptake of TFV and FTC is known, their intracellular accumulation of TFV-DP and FTC-TP, and thus their antiretroviral therapeutic potential, has not been evaluated previously ^15,16^. To address this, we quantified TFV-DP and FTC-TP in astrocytes isolated from brain tissue of uninfected rhesus macaques that received a once daily, 6-month ART regimen that included TFV and FTC (**Figure 1C**). We determined that TFV-DP was quantifiable in astrocytes (15.76 ± 13.92 fmol/million), and that concentrations were similar to reported EC90 for TFV measured in other cell types from rhesus macaques ^17^. In contrast, and consistent with our findings from rhesus macaque brain tissue, FTC-TP was below the limit of quantification in astrocytes (**Figure 1C**). No sex-based differences occurred for astrocyte TFV and FTC quantification (data not shown). We performed similar analyses on brain capillaries, containing brain microvascular endothelial cells and pericytes, from these same macaques. However, cell yields were too low to facilitate detectable quantification (data not shown).

We next performed *in vitro* TFV and FTC treatment for all three cells comprising the BBB as we were unable to perform this comparative analysis with rhesus macaques. These experiments allowed for the determination of whether BBB cells were capable of parent drug metabolism into TFV-DP and FTC-TP. Monocultures of primary human astrocytes, endothelial cells, and pericytes were exposed to a regimen comprised of rhesus macaque equivalent ART doses for 24 hours, after which time the cells were lysed, and TFV-DP and FTC-TP concentrations were determined by LC-MS/MS. In accordance with our previous findings, TFV and FTC did not impact cell viability (data not shown) ^18^. Importantly, all BBB cells expressed cell-type enriched markers present *in vivo*, including VE-cadherin, ANPEP, and GFAP for endothelial cells, pericytes, and astrocytes, respectively, confirming cell identity (**Supplemental Figure 1**). All BBB cell types demonstrated the capability to metabolize TFV and FTC into their active metabolites (**Figure 1D**). However, significant differential concentrations of TFV-DP and FTC-TP occurred among the cells. TFV-DP was 113% (p=0.0337) and 178% (p=0.0220) higher in endothelial cells (6208 ± 109.50 fmol/million) and pericytes (8092 ± 1482 fmol/million), respectively, as compared to astrocytes (2913 ± 493.50 fmol/million) (**Figure 1D**). Similar differential concentrations occurred for FTC-TP, where pericytes (30,723 ± 6220 fmol/million) possessed 804% and 476% higher intracellular levels as compared to endothelial cells (3820 ± 409.60 fmol/million, p=0.0203) and astrocytes (6447 ±1587 fmol/million, p=0.0133), respectively (**Figure 1D**). We also determined that some of the BBB cells exhibited a preferential capacity to metabolize ART. Endothelial cells had 62% higher TFV-DP (6208 ± 109.50 fmol/million) concentrations compared to FTC-TP (3820 ± 409.60 fmol/million, p=0.0094), while the converse occurred in pericytes that had 379% higher FTC-TP (30,723 ± 6220 fmol/million) concentrations than TFV-DP (8092 ± 1482 fmol/million, p=0.0356) (**Figure 1D**). Conversely, there was no statistical difference in the intracellular concentrations of TFV and FTC active metabolites in astrocytes (**Figure 1D**).

### BBB Cells Differentially Express TFV and FTC Transporters and Nucleotide Metabolizing Enzymes

Efflux and influx transporters are well-characterized determinants of brain drug distribution. TFV and FTC are substrates of multiple influx and efflux transporters that may influence the heterogenous ART distribution we observed in **Figure 1**. However, understanding of the differential expression of drug transporters among BBB cell types is limited, as most studies focused solely on endothelial cells ^19,20^. To address this gap, we evaluated TFV and FTC efflux transporters, MRP1 and MRP4, as well as FTC influx transporters, ENT1, OAT1, and OAT3, using proteomics by LC-MS/MS in primary human BBB cell monocultures. Our studies also included primary human hepatocytes as a comparative control as their transporter expression is well characterized (**Figure 2**). Protein expression was also evaluated by Western blot (**Figure 2**).

**Figure 2.**
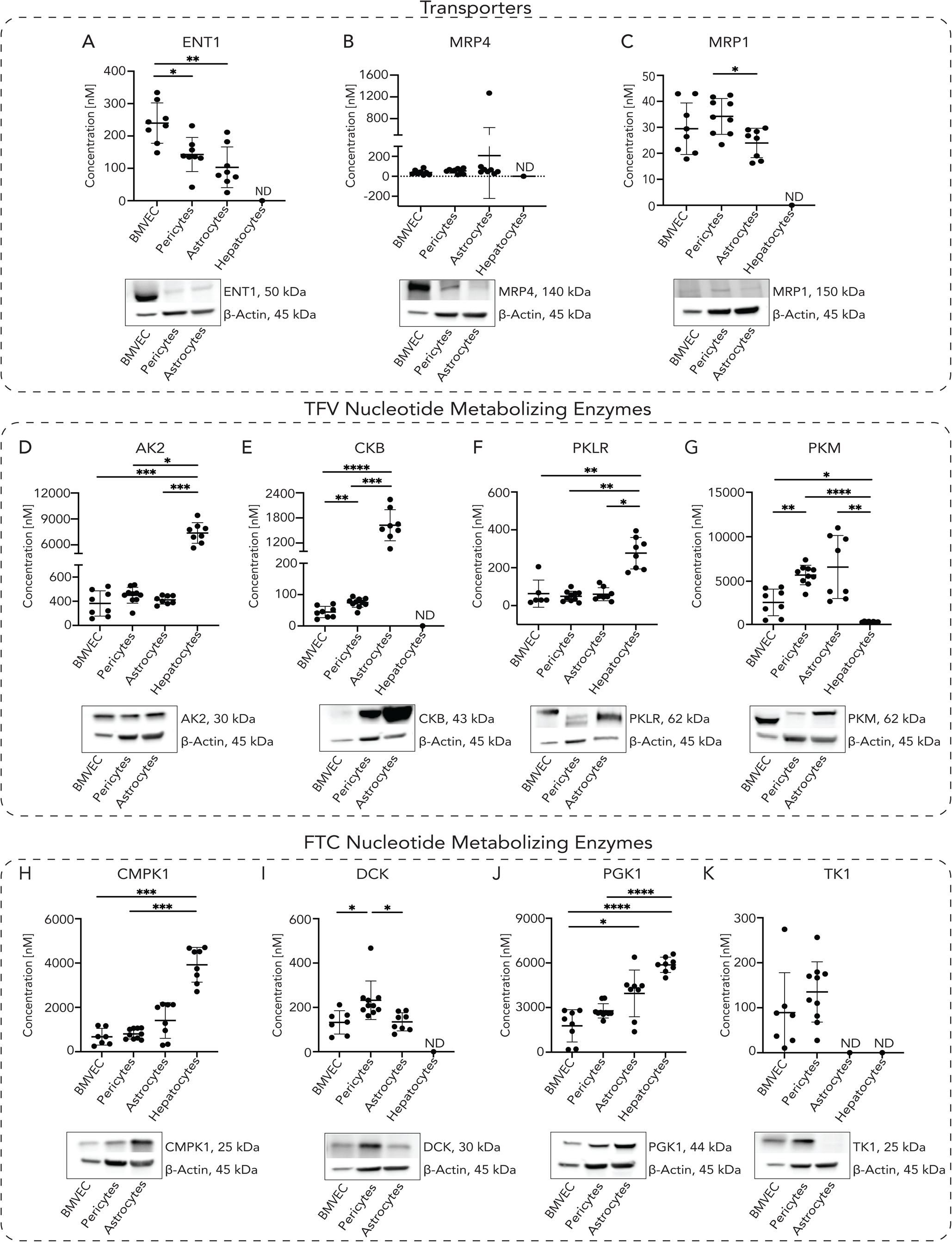
BBB cells differentially express TFV and FTC transporters and nucleotide metabolizing kinases. Primary human BBB cell monoculture and hepatocyte lysates were processed for LC-MS/MS-based proteomics, and protein concentrations were reported by Proteomic Ruler for **(A)** ENT1, **(B)** MRP4, and **(C)** MRP1 TFV/FTC transporters, **(D)** AK2, **(E)** CKB, **(F)** PKLR, and **(G)** PKM TFV nucleotide metabolizing kinases, as well as **(H)** CMPK1, **(I)** DCK, **(J)** PGK1, and **(K)** TK1 FTC nucleotide metabolizing kinases, each with respective protein western blots to confirm protein abundance in BBB cells. Four to five independent experiments with two LC-MS/MS injection replicates each were performed (represented as individual dots). Replicates with missing LC-MS/MS values were omitted from plot. >2 independent experiments with missing values were reported as not reliably detected (ND) by proteomics analyses. Data represented as mean ± standard deviation. Statistical analysis was performed using Brown-Forsythe and Welch ANOVA or Kruskal-Wallis ANOVA. *p < 0.05, **p < 0.01, ***p<0.001, ****p < 0.0001.

We identified differential concentrations of ENT1 and MRP1 among the BBB cell types (**Figure 2A and 2C**), while MRP4 concentrations were consistent (**Figure 2B**). Endothelial cells had significantly higher ENT1 concentrations (239.9 ± 62.36 nM) relative to pericytes (143.0 ± 52.64 nM, p=0.0136) and astrocytes (103.3 ± 62.88 nM, p=0.0019) (**Figure 2A**). However, pericyte MRP1 concentrations (34.22 ± 6.90 nM) were 42% higher compared to astrocytes (23.99 ± 5.65, p=0.0126), while remaining comparable to endothelial cells (29.48 ± 9.90 nM, p=0.6060) (**Figure 2C**). We did not detect TFV influx transporters, OAT1 and OAT3, by LC-MS/MS-based proteomics; however, we previously showed their expression in endothelial cells ^18^. Western blot confirmed transporters expression in all BBB cells. While we did not reliably quantify ENT1, MRP4, and MRP1 in fresh primary human hepatocytes (**Figure 2A, 2B, and 2C**), it is accepted that they are variably expressed in this cell type ^21–23^.

Next, we evaluated the nucleotide metabolizing enzymes that facilitate TFV and FTC metabolism to TFV-DP and FTC-TP. AK2, CKB, PKM, and PKLR were differentially quantifiable in BBB cells (**Figure 2D-2G**). We determined that astrocytes (1626 ± 372.90 nM) expressed CKB approximately 3663% higher relative to endothelial cells (44.39 ± 18.03 nM, p<0.0001) and 2191% higher than pericytes (74.24 ± 14.69 nM, p<0.0001), supporting previous findings (**Figure 2E**) ^24^. Additionally, CKB concentrations were over 67% higher in pericytes (74.24 ± 14.69 nM) relative to endothelial cells (44.39 ± 18.03 nM, p=0.0066) (**Figure 2E**). PKM concentrations were over 224% higher in pericytes (5560 ± 1095 nM) relative to endothelial cells (2482 ± 1500 nM, p=0.0022) (**Figure 2G**). As expected, CKM was not reliably detected in BBB cell types as its expression is limited in the brain ^11^ (data not shown)^10^. Hepatocytes had minimal CKB and PKM concentrations relative to all BBB cell types (**Figure 2E and 2G**).

Similar differential concentrations of FTC metabolizing enzymes also occurred in the BBB cells (**Figure 2H-2K**). Pericytes had greater than 71% DCK concentrations (232.1 ± 87.03 nM) relative to astrocytes (135.0 ± 40.64 nM, p=0.0160) and endothelial cells (132.3 ± 52.88 nM, p=0.0348) (**Figure 2I**). Interestingly, PGK1 was most highly expressed in astrocytes (3951 ± 1569 nM), which had 223% higher concentrations compared to endothelial cells (1771 ± 1085 nM, p=0.0391) (**Figure 2J**). While DCK and TK1 were not reliably detected by LC-MS/MS-based proteomics in hepatocytes, these cells had the highest CMPK1 and PGK1 concentrations relative to BBB cells, supporting their known importance in liver glycolysis (**Figure 2H-2K**) ^25,26^. Western blot confirmed BBB cell nucleotide metabolizing enzyme expression.

### TFV and FTC Transporters are Differentially Active Among BBB cells

Intrigued by the implications of the differential expression of TFV and FTC transporters and nucleotide metabolizing enzymes, we sought to evaluate whether this translated to distinct functional activity. Substrates without overlapping specificity do not exist for ENT1, MRP4, MRP1, OAT1, and OAT3, making it challenging to evaluate their differential activity in our primary BBB cells. Therefore, we evaluated activity of additional efflux transporters known to interact with ART: BCRP, MRP4, and P-gp (**Figure 3**). Importantly, while we did not reliably quantify BCRP and P-gp in our BBB cell types with proteomics by LC-MS/MS, their expression was confirmed with Western blot (**Figure 3D and 3E**).

**Figure 3.**
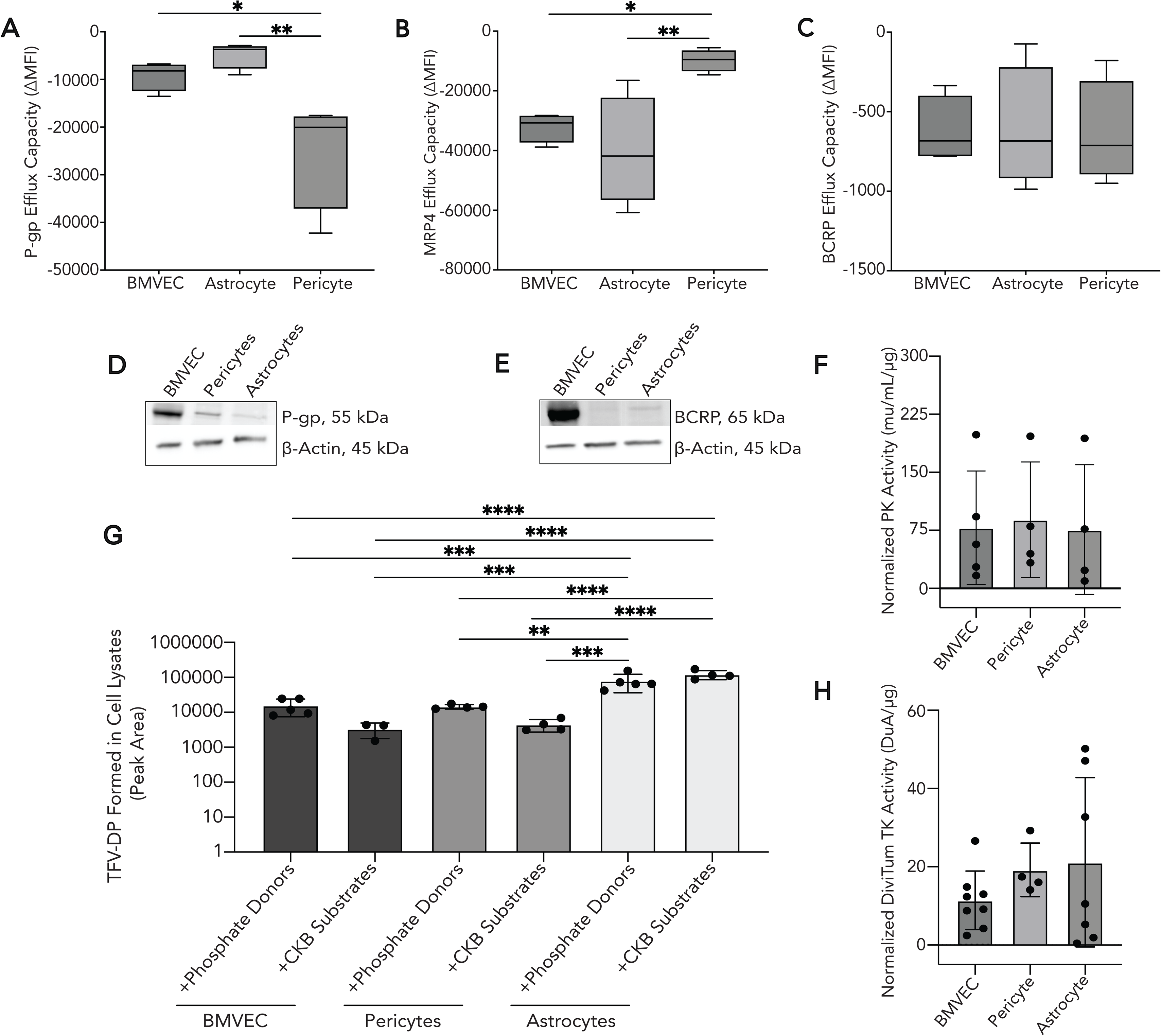
TFV/FTC efflux transporters and nucleotide metabolizing kinases are differentially active in BBB cells. Brain microvascular endothelial cell (BMVEC), pericyte, and astrocyte monocultures were loaded with dyes for **(A)** P-gp (rhodamine 123, 10 μM), **(B)** MRP4 (monobromobimane, 10 μM), and **(C)** BCRP (Hoechst 33342, 5 µg/mL) for 15 minutes at 37°C, 5% CO_2_. The dyes were allowed to efflux out for 2 hours at 37°C, 5% CO_2,_ after which flow cytometry was performed to quantify intracellular fluorescence (mean fluorescence intensity, MFI) as an indicator of efflux capacity. Statistical analysis was performed using one-way ANOVA. **(D)** P-gp and **(E)** BCRP expression were confirmed by western blot, as these proteins were not reliably measurable by proteomics analyses. **(F)** Endogenous pyruvate kinase and **(H)** thymidine kinase activities were measured in primary human BBB cell monocultures by colorimetric activity assay and DiviTum activity assay, respectively. Four to eight independent experiments (represented by individual dots) were performed. Statistical analysis was performed using a Brown-Forsythe and Welch ANOVA test. **(G)** TFV-DP formation in human primary BBB cell lysates was measured by a CKB-mediated tenofovir metabolism assay using LC-MS/MS. Cell lysates were incubated with tenofovir-monophosphate (TFV-MP) and phosphocreatine (+CKB Substrates) or a mixture of phosphocreatine, phosphoenolpyruvate, and ATP (+Phosphate Donors) for 30 minutes at 37°C. The reaction was quenched by LC-MS grade methanol, and peak area of TFV-DP was measured by LC-MS/MS. Three to five independent experiments were performed (represented by individual dots. Statistical analysis was performed using by one-way ANOVA. *p < 0.05, **p < 0.01, ***p<0.001, ****p < 0.0001.

To evaluate efflux transporter activity, primary human BBB cell monocultures were loaded with a fluorescent dye (Monobromobimane for MRP4, Hoechst 33342 for BCRP, and rhodamine 123 for P-gp) whose efflux is known to be mediated by our transporters of interest according to our established method ^18^. We then measured the remaining intracellular fluorescence after sufficient time passed to facilitate dye efflux. Importantly, the fluorescent dyes were not toxic to the BBB cell types (**Supplemental Figure 2C, 2I, and 2O)** and all of the dye rapidly entered the viable cells, as indicated by near 100% positivity for Hoechst 33342, Rhodamine 123, and Monobromobimane (**Supplemental Figure 2D-2F, 2J-2L, 2P-2R**). Efflux of all dyes occurred within two hours, denoted by the significant loss of intracellular fluorescence (**Supplemental Figure 3, black histograms compared to grey histograms**).

We quantified efflux capacity of each transporter by subtracting the difference in intracellular fluorescence, as indicated by the mean fluorescence intensity, following dye uptake and efflux (**Supplemental Figure 4**). We determined that P-gp and MRP4 efflux capacities differed across cell types (**Figure 3A and 3B**). Pericytes had the highest P-gp efflux capacity (−24,671 ± 11,643) as it was 546% and 278% higher than astrocytes (−4520 ± 2823, p=0.0080) and endothelial cells (−8879 ± 3060, p=0.0296), respectively (**Figure 3A**). In contrast, pericytes had the lowest MRP4 efflux capacity (−9815 ± 3722) as it was 75% lower compared to astrocytes (−40,235 ± 18,231, p=0.0095) and 69% lower than endothelial cells (−32,143 ± 4880, p=0.0462) (**Figure 3B**). P-gp and MRP4 efflux activity was comparable between endothelial cells and astrocytes. Surprisingly, BCRP was the only efflux transporter for whom all BBB cells had comparable efflux activity (**Figure 3C**). It is important to note the discordance between efflux expression and activity among the BBB cells, demonstrating the importance of evaluating transporter function.

### Astrocytes Have the Highest CKB Activity, While TK and PK Are Equally Active Among All BBB Cell Types

We first evaluated CKB activity through our TFV activation activity assay, which has been previously used to measure CKB activity in tissues ^10^. In this assay, protein lysates from BBB cell monocultures were incubated for 30 minutes with the TFV-DP precursor, tenofovir monophosphate (TFV-MP), together with phosphocreatine that served as a phosphate donor for the CKB-mediated conversion of TFV-MP to TFV-DP. To measure activity of all TFV-MP metabolizing enzymes, the 1) PKM phosphate donor, phosphoenolpyruvate, 2) universal phosphate donor, ATP and 3) phosphocreatine were incubated with protein from BBB monoculture lysates and TFV-MP. Peak areas of TFV-DP formed from the enzyme-catalyzed reactions were measured in each BBB cell-type for the CKB-mediated conversion (+CKB substrate condition) and general TFV-MP conversion (+Phosphate Donors) (**Figure 3G**). TFV-DP formation in the CKB-mediated reaction was over 3500% higher in astrocytes (120,921 ± 35714) as compared to endothelial cells (3345 ± 1576, p=<0.0001) and pericytes (4439 ± 1744, p=<0.0001) (**Figure 3G**). This also occurred for TFV-DP formation in the general TFV-MP conversion, which was approximately 500% higher in astrocytes (79,229 ± 43,244) relative to endothelial cells (15,616 ± 8159, p=0.0009) and pericytes (14,522 ± 2103, p=0.0016) (**Figure 3G**). Of importance, no significant difference occurred in TFV-DP peak area between the CKB-mediated reactions and the general TFV-MP conversion for all cell types, suggesting that CKB may be the primary enzyme to metabolize TFV-MP to TFV-DP at the BBB as no significant increase in TFV-DP occurred when adding reaction substrates for other TFV activating enzymes. Minimal background from TFV-DP formation was observed in reactions where protein from cell lysate was incubated with TFV-MP without any phosphate donor added, demonstrating assay specificity (data not shown). Further, TFV-DP formation was not detected in negative controls that excluded assay substrates, such as TFV-MP or BBB cell lysates (data not shown).

We next evaluated additional protein kinase activities. We evaluated endogenous pyruvate kinase (PK) function where activity was determined by pyruvate and ATP formation from the PK-catalyzed reaction of phosphoenolpyruvate and ADP. PK activity was comparable across endothelial cells, pericytes, and astrocytes, though enzyme activity varied across experimental replicates (**Figure 3H**). Next, we investigated function of the FTC metabolizing enzyme, thymidine kinase (TK), using a DiviTum^TM^ ELISA assay. TK endogenous activity was comparable among all BBB cell types (**Figure 3H**). However, similar to PK activity, significant variability occurred between measurements, particularly within astrocytes (**Figure 3F**). Our findings demonstrate that while endogenous PK and TK activities are comparably active among BBB cells, their involvement in catalyzing TFV-DP formation occurs in a cell-type specific fashion (**Figure 3F, 3H, and 3G**).

### HIV and ART Regulate TFV and FTC Transporters and Nucleotide Metabolizing Enzymes in Astrocytes and Pericytes, But Not Endothelial Cells

Our study thus far evaluated the presence and activity of transporters and nucleotide metabolizing enzymes at the BBB. However, we focused solely on the basal expression and activity of these proteins, without taking into consideration the potential impact of HIV and ART. There is a substantial impact of HIV, viral proteins, and ART on the BBB; however, most studies focused primarily on BBB permeability and integrity while not considering the components involved in ART transport and metabolism ^27,28^. To evaluate these processes more comprehensively at the BBB in a condition more physiologically relevant to people living with HIV, we evaluated whether HIV and/or ART regulated TFV and FTC transporters and nucleotide metabolizing enzymes. To accomplish this, we treated our primary human BBB cell monocultures with HIV, ART, combined HIV and ART, or vehicle for 24 hours and performed LC-MS/MS-based proteomics. We used HIV and ART concentrations previously measured *in vivo* and commonly used in *in vitro* experiments (**Figure 4**) ^16^.

**Figure 4.**
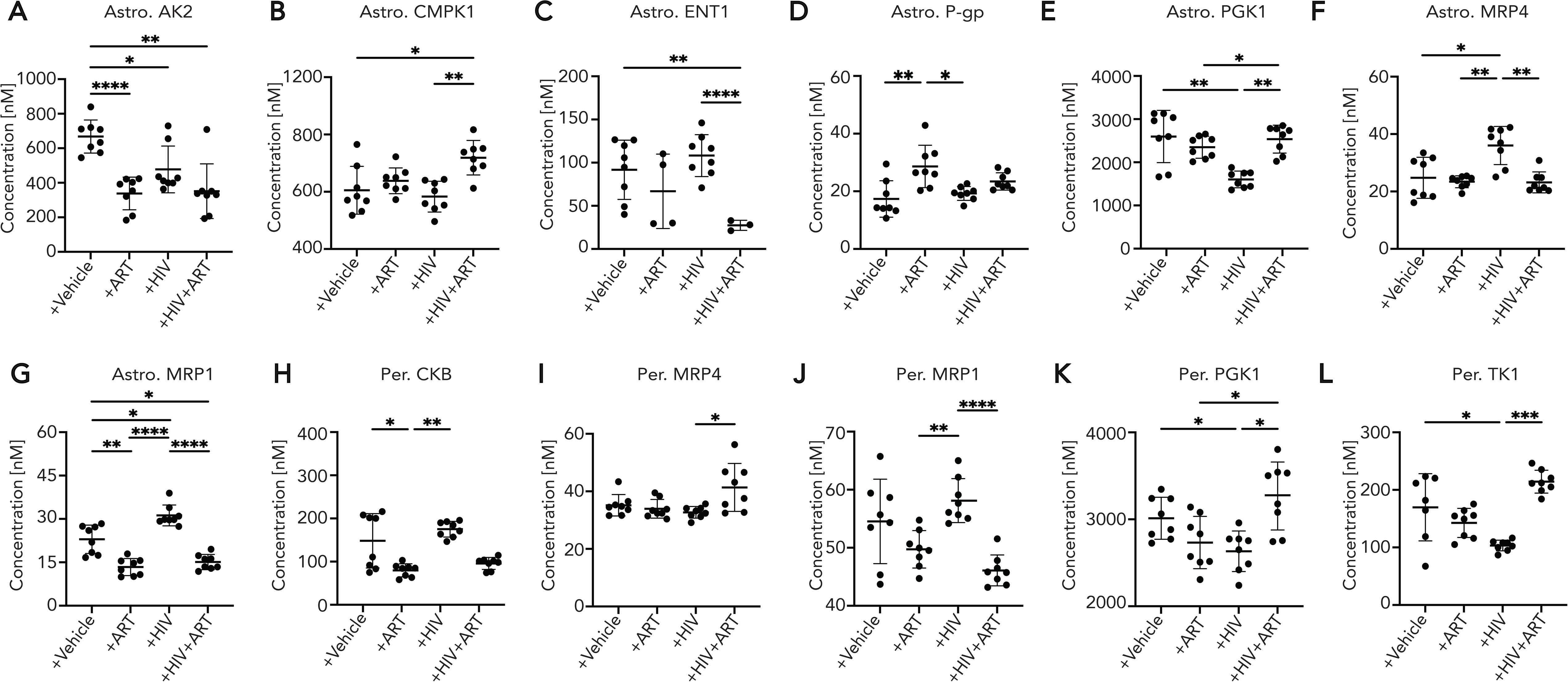
BBB TFV/FTC nucleotide metabolizing kinases and transporters are impacted in a cell-dependent manner by ART and HIV. Primary human pericyte and astrocyte monocultures were exposed to ART (10 μM TFV, 10 μM FTC, and 10 μM DTG), HIV (5 ng/mL), or HIV+ART (10 μM TFV, 10 μM FTC, 10 μM DTG, and 5 ng/mL HIV) for 24 h at 37°C, 5% CO_2_. Treatment with vehicle was used as a control. Pericytes and astrocytes were lysed and processed for proteomics analyses. Concentrations of proteins of interest were measured by Proteomic Ruler. After exposure, changes in protein concentration were assessed in **(A-G)** astrocytes (Astro.) for **(A)** AK2, **(B)** CMPK1, **(C)** ENT1, **(D)** P-gp, **(E)** PGK1, **(F)** MRP4, and **(G)** MRP1. Similarly, changes in protein concentration were assessed in **(H-L)** pericytes (Per.) for **(H)** CKB, **(I)** MRP4, **(J)** MRP1, **(K)** PGK1, **(L)** TK1. Four independent experiments with two LC-MS/MS injection replicates each were performed per condition (represented as individual dots). Replicates with missing LC-MS/MS values were omitted from plot. >2 independent experiments with missing values were reported as not reliably detected (ND) by proteomics analyses. Data represented as mean ± standard deviation. Statistical analysis was performed using Brown-Forsythe and Welch ANOVA or Kruskal-Wallis ANOVA. *p < 0.05, **p < 0.01, ***p<0.001, ****p < 0.0001.

We found that the concentrations of several TFV and FTC transporters and nucleotide metabolizing enzymes were significantly impacted by 24-hour treatment. Strikingly, these changes primarily occurred only in astrocytes and pericytes, but not endothelial cells. HIV, ART, and combined HIV and ART treatment created distinct clustering of pericyte and astrocyte proteomes (**Supplemental Figure 5**). However, this did not occur for endothelial cells as they did not form separate clusters (**Supplemental Figure 5**). This biased impact of HIV and ART was further confirmed as significant changes in TFV and FTC transporter and nucleotide metabolizing enzyme concentrations occurred only in pericytes and astrocytes (**Figure 4**), but not endothelial cells (**Supplemental Figure 6**).

We therefore focused our efforts on understanding the impact of HIV and/or ART on pericytes and astrocytes. ART treatment significantly decreased the concentration of AK2 (338.30 ± 94.74 nM, p=<0.0001) and MRP1 (13.38 ± 2.96 nM, p=0.0028) by ∼50% in astrocytes, as well as CKB (160.30 ± 15.04 nM, p=0.0335) by 30% in pericytes (**Figure 4A, 4G, and 4H**). In contrast, ART increased P-gp concentrations by 65% in astrocytes (28.60 ± 7.35 nM, p=0.0069) (**Figure 4D**). Next, we examined how HIV altered our transporter and metabolizing enzymes of interest. After HIV exposure, TFV and FTC nucleotide metabolizing enzyme concentrations significantly decreased for AK2 (477.70 ± 135.60 nM, p=0.0356) and PGK1 (1522 ± 194.20 nM, p=0.0032) by ∼30% in astrocytes (**Figure 4A and 4E**). In pericytes, HIV decreased PGK1 (2632 ± 233.50 nM, p=0.0356) and TK1 (103.40 ± 9.30 nM, p=0.0409) by 13% and 40%, respectively (**Figure 4K and 4L**). Interestingly, TFV and FTC transporter concentrations increased after HIV exposure for MRP4 (36.02 ± 6.63 nM, p=0.0314) by 45% and by 35% for MRP1 (31.22 ± 3.61 nM, p=0.0117) in astrocytes, but there were no impacts on TFV/FTC transporter concentrations in pericytes (**Figure 4F, 4G, 4I, and 4J**).

Finally, we determined the combined impact of HIV and ART. Combined HIV and ART treatment decreased AK2 and MRP1 concentrations relative to vehicle only for astrocytes but not pericytes (**Figure 4A and 4G**). Interestingly, astrocyte exposure to the combination of HIV and ART resulted in a synergistic effect for CMPK1 and ENT1, where the concentrations of these proteins changed only upon dual treatment but not with their individual exposure (**Figure 4B and 4C**). Astrocyte CMPK1 concentrations increased by 18% (719.0 ± 60.48 nM, p=0.0448), while ENT1 concentrations decreased by 70% (27.41 ± 5.89 nM, p=0.0050) (**Figure 4B and 4C**). In contrast, neither of these proteins were changed in pericytes with any treatment condition (data not shown). Taken together, these data indicate that HIV, ART, and combination treatment may profoundly impact the transport and metabolic capacity of the BBB, which occurs by regulating pericytes and astrocytes, but not endothelial cells (**Supplemental Figure 6**). Further, there is a protein-specific manner for which exposure to HIV, ART, or a combination of HIV and ART selectively regulates proteins involved in TFV and FTC transport and metabolism.

### HIV and ART Modulate Astrocyte and Pericyte Cell Transport and Regulation, Metabolism, Immune Response, and Cell-Cell Communication Pathways

While we focused primarily on TFV and FTC transporters and metabolizing enzymes, we also recognize that HIV and ART impact additional cellular functions at the BBB. So, we next evaluated pharmacologically related cellular pathways that may contribute to the protein concentration changes we observed (**Figure 4**) following HIV, ART, and combination treatment. We used SimpliFi^TM^ to accomplish this by further processing and analyzing BBB proteomes to identify additional cellular pathways of interest. We identified several proteins as significantly impacted by HIV or ART exposure (p<0.05), relative to vehicle, that can be publicly <u>accessed</u>. To summarize and organize these findings, we further filtered differentially expressed proteins such that the log2 fold change within affected pathways of interest relative to vehicle was greater than 0. We found that the resulting top hits in both astrocytes and pericytes (p<0.05, log2 fold change>0) were pathways involved in 1) cell transport and regulation, 2) metabolism, 3) immune response and signaling, 4) cell-cell communication, and 5) HIV pathogenesis (**Figure 5**). Astrocytes are actively infected by HIV, and thus, we were not surprised to identify several pathways implicated in immune responses and signaling and HIV pathogenesis in HIV-exposed astrocytes (**Figure 5A and 5C**) ^29^. Pericytes are also permissive to HIV and we observed pathway hits involved in host-interactions for HIV pathogenesis and immune response and signaling after exposure to HIV, ART, and a combination of HIV and ART, which were characterized by a mixture of proteins that were both upregulated and downregulated (**Figure 5D-5F**) ^30^. Similar to that which occurred for TFV and FTC transport and metabolizing proteins, endothelial cell exposure to HIV, ART, and combined HIV and ART treatment had minimal impact in these pathways (**Supplemental Figure 6K, 6L, and 6M**).

**Figure 5.**
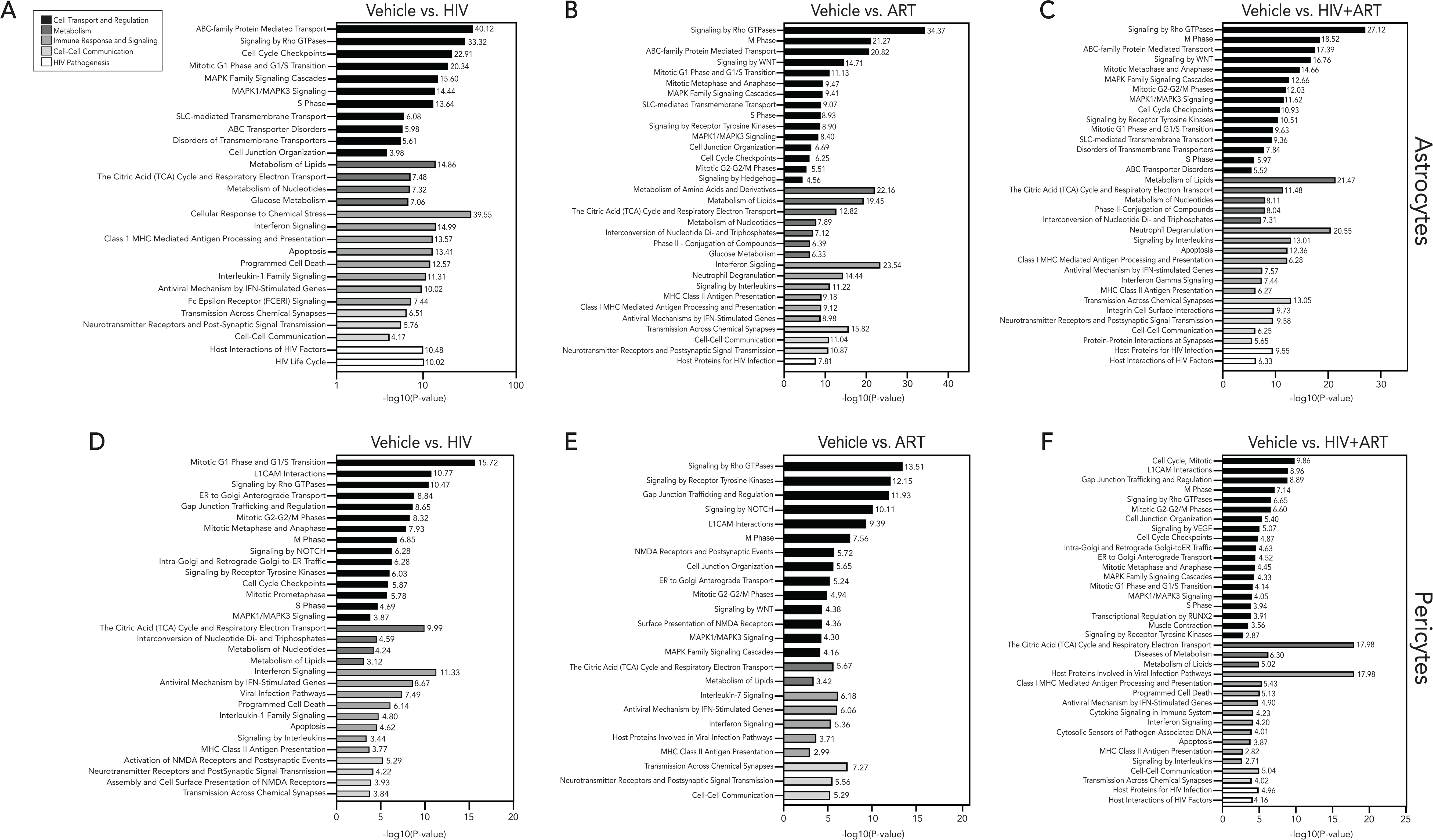
HIV and ART differentially impact astrocyte and pericyte cell pathways involved in cell transport and regulation, metabolism, immune response and signaling, and cell-cell communication. Primary human pericyte and astrocyte monocultures were exposed to ART (10 μM TFV, 10 μM FTC, and 10 μM DTG), HIV (5 ng/mL), or HIV+ART (10 μM TFV, 10 μM FTC, 10 μM DTG, and 5 ng/mL HIV) for 24 h at 37 °C, 5% CO_2_. Treatment with vehicle was used as a control. Pericytes and astrocytes were lysed and processed for proteomics analyses. Pathways significantly altered in astrocytes after **(A)** HIV, **(B)** ART, and **(C)** HIV+ART exposure as well as in pericytes after **(D)** HIV, **(E)** ART, and **(F)** HIV+ART (relative to vehicle) were analyzed by SimpliFi proteomics software. Pathways of interest were filtered by pathway changes greater than one log fold and hypergeometric p-value < 0.05. Pathway hits of interest were ranked by –log10(hypergeometric p-value). Pathway analyses reflect four independent experiments per condition with two LC-MS/MS injection replicates each.

## Discussion

The BBB represents a significant obstacle in CNS drug penetration that challenges efforts in treatment of neurological disorders. This emphasizes the critical importance of understanding the molecular mechanisms that govern CNS drug disposition at the BBB. Here, we use first-line ART drugs, TFV and FTC, as models to investigate the mechanisms of drug transport and metabolism at the BBB that may influence CNS drug access. Our study demonstrates that the BBB is a pharmacologically dynamic microenvironment that possesses differential activities of drug metabolizing enzymes and transporters within endothelial cells, astrocytes, and pericytes, which are distinctly impacted by disease and therapeutic exposure. Surprisingly, our findings identify pericytes and astrocytes as underappreciated contributors to therapeutic access to the brain. Further, our work highlights pharmacologically active pathways at the BBB that may regulate CNS drug disposition and impact therapeutic efforts to alleviate neurologic disease.

Evidence suggests ART distributes heterogeneously across brain regions ^8,9,31,32^, though this is an emerging area of investigation. To contribute further to this knowledge, we measured TFV, FTC, and their metabolites, TFV-DP and FTC-TP, respectively, across three brain regions from the well-characterized SIV-infected rhesus macaque model. Our findings demonstrate that, overall, TFV had greater CNS access than FTC. This is striking as these therapies belong to the same ART class and have similar molecular profiles, including charge and molecular weight. Of importance, these *in vivo* findings confirm our previous work using an *in vitro* model of the human BBB indicating the relevance of this model as a tool in evaluating CNS ART disposition ^18^. Our current work confirms that variable concentrations of TFV and FTC occured among brain regions, although no sex-based differences occurred. Further, we showed, for the first time, that this regional variability extended to TFV and FTC active metabolites. We identified the highest TFV and TFV-DP concentrations in the cerebellum, which is vulnerable to HIV infection and damage ^33^. These findings are intriguing as, historically, CSF measurements used to predict CNS ART suggested increased FTC penetrance in brain relative to TFV, which our findings do not support ^34,35^. Rather, our results further highlight the disparity between CSF and brain tissue ART measurements and caution against its use as a proxy for parenchymal concentrations ^34,35^. Our findings are consistent with the ART heterogeneity observed in human and mouse brains, but are distinct from other rhesus macaque studies ^7,36^, that may be attributed to differences in study design, including viral strain, ART regimen, and dosing duration. Our results continue to elucidate the differential distribution of TFV and FTC within the CNS and, more importantly, the heterogenous measurement of their pharmacologically active metabolites across brain regions.

Drug transport is most widely studied in the context of peripheral organs, primarily liver. As the proteins involved in this process are also present at the BBB, it has been assumed that the same mechanisms extend to the brain. However, it is unlikely that transport mechanisms are identical between the CNS and peripheral organs due to the specialized environment required for neuronal survival. An additional challenge has been the focus of evaluating drug transporters in endothelial cells, some of which have not been CNS-derived, to the exclusion of pericytes and astroytes ^19,20,37^. Our work determined that, while present, each BBB cell type expressed differentially active TFV and FTC transporters. Surprisingly, we demonstrated that pericytes and astrocytes expressed several functionally active TFV and FTC efflux and influx transporters. Often, these cells had significantly higher transporter expression than endothelial cells, which may explain their differential intracellular TFV and FTC accumulation ^15,16,18^. To evaluate the transport potential of the BBB relative to liver, we compared drug transporters in primary human hepatocytes relative to our BBB cells. We found little hepatocyte ENT1, MRP4, and MRP1, which were prevalent in BBB cells. Previous reports suggest ENT1 and MRP4 are highly variable in hepatocytes, whereas MRP1 is low, likely explaining our inability to reliably quantify these transporters by mass spectrometry-based proteomics ^22,38,39^. These findings demonstrate the importance of evaluating drug transport at the BBB, rather than viewing it as an extension of the liver, as it possesses distinct mechanisms that may contribute to differential drug disposition in the CNS. Additionally, our results elucidate the importance in considering pericytes and astrocytes, in addition to brain endothelial cells, in establishing transport function at the BBB.

Prior studies observed that TFV and FTC accumulate intracellularly within endothelial cells, pericytes, and astrocytes that results in differential cell-type specific concentrations ^15,16^. Although these findings suggest that BBB cells may have distinct capabilities in accumulating TFV and FTC, prior studies did not evaluate whether BBB cells were capable of TFV and FTC metabolism. Using astrocytes isolated from the brains of healthy rhesus macaques dosed with ART once daily for 6 months, we reported variable concentrations of TFV-DP that were similar to TFV EC90s previously determined in other macaque cell types ^17^. These measurements may indicate ART therapeutic potential in astrocytes, which is essential to viral eradication of the brain as these cells are infected by HIV and contribute to a persistent CNS reservoir ^29,40^. We did not detect FTC-TP in rhesus macaque astrocytes, and were surprised to quantify FTC-TP in human astrocytes *in vitro*. This finding suggests that astrocytes possess the capacity to pharmacologically transform FTC into its active metabolite, should efficacious doses be achieved in the brain. We did not have sufficient cell yields to quantify active ART metabolites in rhesus macaque capillaries, however our *in vitro* treatments demonstrated that endothelial cells and pericytes also had the capacity to form active TFV and FTC metabolites. While the metabolism of TFV and FTC in BBB cells initially appears promising for treating the brain during HIV, our findings raise concern for the utility of nucleotide analogues as putative CNS therapies. Our work suggests that local drug metabolism in endothelial cells, pericytes, and astrocytes may limit drug availability in the brain parenchyma by restricting it to cells at the BBB, as to our knowledge, there are no known transporters for the negatively charged, high molecular weight, and bulky TFV and FTC metabolites – suggesting they may be retained within these cells rather than being released into the brain parenchyma.

Here, we determined that pericyte and astrocyte TFV and FTC nucleotide metabolizing enzymes are comparable to, or significantly higher than, endothelial cells. For many nucleotide metabolizing enzymes, BBB cell concentrations were comparable to, or higher, than hepatocytes indicating that these cells hold competitive ART metabolizing capabilities as the liver. This was intriguing as the brain is not typically regarded as a drug metabolizing organ. Taken together, these results highlight that BBB cells harbor the biotransformation machinery necessary to differentially metabolize TFV and FTC, identifying it is a pharmacologically relevant microenvironment wherein drug metabolism is a pertinent mechanism in determining CNS drug availability.

The BBB responds dynamically to disease that may lead to dysregulation, exacerbating neurologic injury and dysfunction. We determined that BBB cells were differentially impacted by HIV and ART by promoting changes in cell pathways related to transport, mitochondrial dysfunction, oxidative stress, immune regulation and response, and inflammation. Interestingly, alterations in transporter and nucleotide metabolizing enzyme expression were restricted to pericytes and astrocytes. This was unexpected as previous studies reported changes in brain endothelial cell pathways, such as transport, after exposure to ART or the HIV protein tat ^28,41,42^. The disparity in our findings may occurr due to our use of clinically relevant concentrations of ART and using replication competent HIV, rather than a viral protein. We propose evaluation of CNS drug disposition occur at baseline levels and also in the context of the diseased state, as the fundamental physiology of the BBB, including drug transporters and metabolizing enzymes, differs during pathology, effectively shaping CNS drug availability.

In summary, first-line ART drugs, TFV and FTC, were evaluated as representative models of CNS-penetrant small molecule drugs with significant disease relevance to determine mechanisms at the BBB that may govern heterogenous CNS drug disposition. We establish a foundation regarding BBB cells as pharmacological entities that form the dynamic BBB microenvironment and may determine CNS drug access as they possess machinery to differentially transport and metabolize CNS-penetrant drugs. Our findings provide evidence of dynamic pharmacological capabilities of metabolism and transport within BBB cells and provide further considerations of region-specific differences in CNS drug metabolism, transport, and distribution. Additionally, our results highlight that mechanisms of drug transport and metabolism at the BBB may be impacted by therapeutic intervention and disease, which may influence CNS drug disposition.

### Limitations of the study

First, quantification of parent TFV and FTC and their metabolites occurred in a small number of animals, which limits determinations of sex-based differences. Second, transporter and metabolizing enzymes concentrations produced by the Proteomic Ruler are based on approximate estimations of cell ploidy, total cellular protein concentration, and cell size determined by previous studies ^43,44^. Additionally, demographic information such as age, sex, gender, and health status was limited for primary human BBB cells and hepatocytes used in this study. Finally, while concentrations of HIV and ART were used according to previous *in vitro* studies and *in vivo* measurements, we cannot fully predict the extent of metabolite formation and cell pathway impacts at the BBB *in vivo* using cell monocultures alone, which will require further studies. Taken together, while limitations exist within our study, it establishes a strong foundation for the investigation of differential cell capabilities for ART metabolism and transport at the BBB.

## Supporting information

Key Resources Table

## Acknowledgements

We thank Mr. Mark Marzinke of the Clinical Pharmacology Analytical Lab for his assistance with antiretroviral therapy determination. We are grateful to Tarsh Shah for his assistance in optimizing the initial proteomics workflow for pericytes and astrocytes exposed to HIV or ART. We also thank Alejandro J. Brenes at the University of Dundee for guidance on the use of the Proteomic Ruler for our applications used in this study. This research was funded by the National Institutes of Health under award number R00 DA044838 (DWW), R01 DA052859 (DWW), and U01 DA058527 (DWW), R01 GM103853 (BCO), and R01 AG064908 (BCO). Additionally, HNW was supported by T32 GM144272 granted to the Biochemistry, Cellular & Molecular Biology Graduate Program at Johns Hopkins. This work was supported, in part, by the Center for AIDS Research at Emory University (P30 AI050409). The content is solely the responsibility of the authors and does not necessarily represent the official views of the National Institutes of Health.

## Author contributions

H.N.W., B.C.O., and D.W.W. conceived and designed the study. H.N.W., S.K., and K.H. performed the experiments. H.N.W. and A.W. created the accessible cell repository. H.N.W. and D.W.W performed the data curation and data analysis. H.N.W., B.C.O., R.C.O., and D.W.W. drafted and edited the manuscript.

## Supplemental Information

Document S1. Figures S1-S6

## STAR METHODS

### Resource availability Lead contact

Further information and requests for resources and reagents should be directed to and will be fulfilled by the Lead Contact, Dionna Williams.

### Materials availability

We did not generate new unique reagents in this study.

### Data and code availability

[MassIVE](https://massive.ucsd.edu/ProteoSAFe/private-dataset.jsp?task=5846f3074fbe43418a527d3b6e81d26f); Password: WilkinsBBB

[SimpliFi](https://simplifi.protifi.com/#/p/55ab4180-da36-11ee-a4d0-6990211fd4ff)

BBB Repository Code (shinyApps)

Any additional information required to reanalyze the data reported in this paper is available from the lead contact upon request.

### Experimental Model and Subject Details

#### Cell lines

Primary human astrocytes (ScienCell Research Laboratories, Carlsbad, CA) were grown to confluence in Basal Medium Eagle (Thermo Fisher Scientific, Waltham, MA) buffered to pH 7.2–7.5 with 2.2 g/L sodium bicarbonate and 15 mM HEPES (Gibco, Grand Island, New York). Media was supplemented with 2% fetal bovine serum (FBS) (R&D Systems, Minneapolis, MN), 1% penicillin–streptomycin 10,000U/mL (Gibco), and 1% astrocyte growth supplement (ScienCell Research Laboratories). Astrocytes were used at passages 1–6 for all experiments. Information on donor including sex and age were not available from source.

Primary human brain microvascular endothelial cells (Cell Systems, Kirkland, WA) were grown to confluence on tissue culture plates coated with 0.2% gelatin (Thermo Fisher Scientific) in medium 199 (M199) (Gibco) buffered to pH 7.2–7.5 with 2.2 g/L sodium bicarbonate and 15 mM HEPES (Gibco). Complete M199 media (M199C) was supplemented with 20% heat-inactivated newborn calf serum (Gibco), 1% penicillin–streptomycin 10,000U/mL (Gibco), 25 mg/L heparin (Sigma, St. Louis, MO), 5% heat-inactivated human serum AB (GeminiBio, Sacramento, CA), 50 mg/L ascorbic acid (Sigma), 7.5 mg/L endothelial cell growth supplement (Sigma), 2 mM L-glutamine (Gibco), and 5 mg/L bovine brain extract (Lonza, San Diego, CA). Endothelial cells were used at passages 2–12 for all experiments. Information on donor including sex and age were not available from source.

Primary human pericytes (ScienCell Research Laboratories) were grown to confluence on 1.5% poly-L-lysine coated flasks (Sigma) in Pericyte Medium (ScienCell Research Laboratories), containing HEPES and sodium bicarbonate buffered to pH 7.4. Media was supplemented with 2% fetal bovine serum (FBS) (ScienCell Research Laboratories), 1% penicillin-streptomycin 10,000U/mL (Gibco), and 1% pericyte growth supplement (ScienCell Research Laboratories). Pericytes were used at passages 2-13 for all experiments. Information on donor including sex and age were not available from source.

Fresh primary human hepatocytes (BioIVT, Westbury, NY) were purchased in liquid suspension and isolated from a deceased Caucasian female, age 66 with a healthy medical background. Gender information was not available from source.

The manufacturers of the primary human brain microvascular endothelial cells, pericytes, and astrocytes confirmed the cellular origin, functional capacity, and HIV and mycoplasma status prior to distribution. Upon receipt, all primary human brain endothelial cells, pericytes, astrocytes, and hepatocytes were authenticated by immunofluorescence or mass spectrometry-based proteomics for cell-type enriched markers.

To isolate rhesus macaque astrocytes, three female and two male rhesus macaque whole brain sections were washed in isolation media containing Hank’s Balanced Salt Solution (HBSS) (Thermo Fisher Scientific) followed by isolation media containing Dulbecco’s Modified Eagle Medium (DMEM) (Thermo Fisher Scientific), 1% HEPES (Gibco), 1% L-glutamine (Gibco), 0.04% geneticin (Thermo Fisher Scientific), 1% penicillin–streptomycin 10,000U/mL (Gibco), and 100mM sodium pyruvate (Thermo Fisher Scientific). Blood vessels were surgically removed from whole brain sections, and brain tissue was subsequently washed with PBS buffered to pH 7.4 (Thermo Fisher Scientific) and digested for 30 minutes at 37°C, 5% CO_2_ with a trypsin solution containing 10% of 2.5% trypsin (Thermo Fisher Scientific), Dulbecco’s Modified Eagle Medium (Thermo Fisher Scientific), DNase at a concentration between 50 to 375 U/μl (Thermo Fisher Scientific), and 1% geneticin (Thermo Fisher Scientific). Digested brain tissue was filtered twice using 183 μm sterile mesh (Corning, Corning NY) and a 100 μm nylon cell strainer (Corning) in 10% DMEM (Thermo Fisher Scientific) and centrifuged to isolate cells. Astrocytes were then isolated using a density gradient containing percoll (Thermo Fisher Scientific) and 1X phosphate buffered saline (Gibco), filtered through 40 μm nylon cell strainer (Corning), centrifuged, and washed with DMEM-10 supplemented with 15% fetal bovine serum (Thermo Fisher Scientific), 0.1% geneticin (Thermo Fisher Scientific), and 100 mM sodium pyruvate. Isolated astrocytes were counted, pelleted, and lysed in 70% LC-MS-grade methanol. Proteomics by mass spectrometry was used to authenticate isolated astrocytes for cell-type enriched markers.

Brain capillaries were isolated concurrent with astrocyte isolation, wherein digested macaque brain tissue was filtered twice using 183 μm sterile mesh (Corning) and a 100 μm nylon cell strainer (Corning) in 10% DMEM (Thermo Fisher Scientific). The resulting brain capillaries were removed from the 100 μm nylon cell strainer (Corning) and centrifuged, pelleted, and lysed in 70% LC-MS-grade methanol for mass spectrometry analyses.

#### Viruses

HIV-1_ADA_ (ARP-416) was obtained through the NIH HIV Reagent Program, Division of AIDS, NIAH, NIH: Human Immunodeficiency Virus-1 ADA, ARP-416 contributed by Dr. Howard Gendelman. HIV-1 p24 protein was quantified by HIV-1 p24 ELISA (Perkin Elmer, Shelton, CT) according to manufacturer’s instructions and subsequently aliquoted. An aliquot of the original SIVmac251 viral stock produced by the laboratory of Ronald Desrosiers was obtained for macaque inoculation and expanded by infecting rhesus macaque peripheral blood mononuclear cells (PBMCs). Further SIVmac251 processing and evaluation was conducted as described previously^45^.

#### Ethics and biosafety statement

Rhesus macaques (*macaca mulata*) originating from The Johns Hopkins University School of Medicine rhesus macaque colony were used in this study. Rhesus macaques were selected by pre-screening negative for the immunoprotective MHC class I alleles *Mamu-A*01*, *Mamu-B*08*, and *Mamu-B*17*. All rhesus macaques were approximately 6 kg throughout the study and when tissues were harvested.

| Macaque ID | Sex | Developmental Stage | Age (years) |
| --- | --- | --- | --- |
| M1 | M | Juvenile | 4 |
| M2 | M | Juvenile | 4 |
| M3 | M | Juvenile | 4 |
| F1 | F | Juvenile | 4 |
| M4 | M | Juvenile | 4 |
| F2 | F | Juvenile | 4 |
| M5 | M | Juvenile | 4 |
| F3 | F | Juvenile | 4 |

All procedures were performed in compliance with Animal Care and Use Committee and performed at the Johns Hopkins University and Emory University according to the guidelines of the US Department of Agriculture Animal Welfare Act and the NIH Guide for the Care and Use of Laboratory Animals. All protocols were carried out following the guidelines of The Johns Hopkins Medical Center and Emory University School of Medicine.

#### Animals and study design

We used a well-established rhesus macaque (*macaca mulatta*) model optimal for studying the impact of therapeutics on HIV outcome ^7,36,46^. The present study was conducted using macaque samples from two cohorts. The first cohort involved two SIV-infected, ART-treated rhesus macaques (M1 and M2) and one healthy, ART untreated rhesus macaque (M3) that were inoculated intravenously using SIVmac251 as previously described ^45^. At day 42 post infection, a subcutaneous dose of 2.5 mg/kg dolutegravir (DTG, Viiv, London, England, UK), 20 mg/kg tenofovir (TFV, Gilead, Foster City, CA), and 40 mg/kg emtricitabine (FTC, Gilead) was administered once daily for six months. Euthanasia occurred at approximately 200 days post-inoculation. The second cohort involved five uninfected, ART-treated rhesus macaques (M4-M5, F1-F3). ART administration and euthanasia occurred in the same manner as described for cohort one animals, except an intramuscular dose of 2.5 mg/kg dolutegravir (DTG, Viiv, London, England, UK), 5.1 mg/kg tenofovir, and 50 mg/kg emtricitabine (FTC, Gilead) was used.

### Method details

#### Immunofluorescent microscopy

Primary brain endothelial cells, pericytes, and astrocytes were seeded (1.6 x 10^4^ cells/dish) on 35 mm ibiTreat dishes (Ibidi USA, Madison, WI). Brain endothelial cells were seeded on ibiTreat dishes coated with 0.2% gelatin. Upon growth to 80% confluence, cells were fixed with 4% paraformaldehyde (Electron Microscopy Sciences, Hatfield, PA) for 15 minutes. The cells were stained with wheat germ agglutinin conjugated to Texas red (Thermo Fisher Scientific), for 10 min to facilitate identification of cell morphology through staining of the plasma membrane. Cells were then permeabilized in 0.01% Triton X-100 (Sigma) for one minute then blocked for two hours at room temperature in Dulbecco’s phosphate-buffered saline without calcium or magnesium (DPBS) (Thermo Fisher Scientific) containing 5 mM ethylenediaminetetraacetic acid (EDTA) (Sigma), 1% fish gelatin (Sigma), 1% essentially immunoglobulin-free bovine serum albumin (Sigma), 1% heat-inactivated human serum AB (GeminiBio, Sacramento, CA), and 1% goat serum (Vector Laboratories, Newark, CA). Cells were probed with antibodies with specificity to GFAP, VE-Cadherin, or ANPEP overnight at 4°C, washed three times with DPBS at room temperature, and probed with the appropriate Alexa Fluor 488 conjugated secondary antibody for one hour at room temperature. Immunofluorescence microscopy was performed as previously described ^18^. In short, the ibiTreat plates were stained with Ibidi mounting medium containing DAPI as a counterstain to identify nuclei (Ibidi). All antibodies were titered to determine optimal concentrations. Antibody details are listed in the Key Resources Table. Isotype-matched controls, staining with only secondary antibodies, and unstained cells were used as negative controls and to account for autofluorescence and nonspecific signal from the cells. The ibiTreat plates were imaged by fluorescent microscopy using the ECHO Revolution (San Diego, CA). Images were acquired 1–15 days post fixation using the ECHO Revolution, as previously ^18^. Fiji v2.14.0 (National Institutes of Health) was used for image analysis.

#### ART Intracellular Concentration Determination

Primary human brain endothelial cells were seeded on 10 cm plates at a density of 1 x 10^6^ cells/plate. Primary human pericytes were seeded on 150 cm^2^ flasks at a density of 1 x 10^6^ cells/flask. Primary human astrocytes were seeded in 6-well plates at a seeding density of 0.3 x 10^6^ cells/well. Upon reaching 90% confluency, cells were treated with TFV (18 µM), FTC (2 µM), DTG (6 µM), or vehicle, for 24 hours at 37°C, 5% CO_2_ (TFV, FTC, and DTG were all from Toronto Research Chemicals, Toronto, Canada). TFV, TFV and FTC were prepared in a 100 mM stock solution in sterile water, while DTG was diluted in DMSO together with sterile water. All subsequent dilutions of DTG were made in sterile water, and the final concentration of DMSO was less than 0.1%. Of note, ART concentrations reflect the approximate equivalent concentrations of the macaque ART dosing regimens used in this study.

*In vitro* ART-treated brain endothelial cells, pericytes, and astrocytes, as well as rhesus macaque brain tissue and isolated astrocytes, were lysed in 70% LC-MS-grade methanol. FTC, TFV, FTC-TP, and TFV-DP were quantified in collaboration with the Clinical Pharmacology Analytic Laboratory at The Johns Hopkins University School of Medicine as previously described ^47,48^. Briefly, FTC and TFV were analyzed in positive mode using a TSQ Vantage® triple quadrupole mass spectrometer coupled with a HESI II® probe (Thermo Scientific). The analytical run time was 8 min, and the assay lower limits of quantitation were 5 fmol/sample and 50 fmol/sample for TFV-DP and FTC-TP, respectively. The lower limits of quantification for TFV and FTC were 0.05 ng/sample and 0.25 ng/sample, respectively. Each experimental replicate of endothelial cells, pericytes, and astrocytes includes data collected from three to four independent experiments. Experimental replicates for astrocytes isolated from rhesus macaques were collected from five independent animals. Lastly, each experimental replicate for ART-treated rhesus macaque brain includes data collected from two independent animals.

#### Western Blot

Endothelial cells, pericytes, and astrocytes were lysed with 1X RIPA buffer (Cell Signaling Technology, Danvers, MA) supplemented with 1X protease/phosphatase inhibitor (Cell Signaling Technology). Total protein concentrations were determined by Bradford Assay with the Bio-Rad Protein Assay Dye reagent concentrate (Bio-Rad, Hercules, CA) following the manufacturer’s instructions. 40 μg of protein was electrophoresed on a 4–12% polyacrylamide gel (Bio-Rad) and transferred to nitrocellulose membranes (Amersham Biosciences, Woburn, MA). Membranes were blocked for two hours at room temperature with 5% nonfat dry milk (Lab Scientific bioKEMIX Inc., Danvers, MA) and 3% bovine serum albumin (Thermo Fisher Scientific) in 1X Tris-Buffered Saline (Quality Biological, Gaithersburg, MD) containing 0.1% Tween-20 (TBS-T, Sigma). Blots were probed with antibodies with specificity to BCRP, P-gp, MRP4, MRP1, ENT1, PGK1, AK2, CKB/CKM, TK, CMPK, DCK, PKM1, and PKLR overnight at 4°C, washed with TBS-T, and probed with the appropriate secondary antibody for one hour at room temperature. Antibody details are provided in the Key Resources Table. All antibodies were used at concentrations recommended by manufacturer. Western Lightning Plus-ECL (PerkinElmer, Waltham, MA) was used as chemiluminescence substrate, and the signal detected with the Azure Biosystems c600 Imager (Azure Biosystems, Dublin, CA). As a loading control, membranes were stripped with Restore Plus Western Blot Stripping Buffer (Thermo Fisher Scientific) and re-probed with antibody against β-Actin HRP for one hour at room temperature.

#### Blood brain barrier monoculture proteomics by LC-MS/MS

Endothelial cells, pericytes, and astrocytes were lysed with a 1X proteomics lysis buffer 5% sodium dodecyl sulfate (SDS) (Invitrogen, Carlsbad, CA) and 50 mM triethylammonium bicarbonate (Sigma). Protein concentration was determined using the Pierce BCA Protein Assay Kit (Thermo Fisher Scientific) following the manufacturer’s instructions. Proteins were digested into peptides using the S-Trap™ micro spin columns (ProtiFi, Fairport, NY) following the manufacturer’s instructions. In brief, 100 μg of protein from each sample was solubilized in 5% SDS, reduced with 120 mM tris(2-carboxyethyl)phosphine (ProtiFi) for 15 minutes at 55°C, alkylated using 500 mM methyl methanethiosulfonate (ProtiFi) for 10 minutes at room temperature acidified with 1.2% phosphoric acid (ProtiFi), trapped on column, and then digested by 20 μg MS-grade trypsin (Thermo Fisher Scientific). Once peptides were eluted, they were dried under vacuum centrifugation (Eppendorf, Enfield CT) overnight, resuspended in 300 μl of 0.1% formic acid in water (v/v) and de-salted according to the manufacturer’s instructions for the Pierce^TM^ peptide desalting spin columns (Thermo Fisher Scientific). Once peptides were desalted, they were dried under vacuum centrifugation (Eppendorf, Enfield, CT) overnight, resuspended in 100 μl 0.1% formic acid in H_2_O (Thermo Fisher Scientific), and quantified according to the manufacturer’s instructions using the Pierce Quantitative Colorimetric Peptide assay (Thermo Scientific). Samples were diluted to 100 ng/μl and 2 μl were injected by an EasyNLC 1200 (Thermo Fisher Scientific) nanoflow liquid chromatography system coupled to a timsTOF FleX mass spectrometer (Bruker, Billerica, MA). Mobile phase A was 0.1% formic acid in H_2_O (Thermo Fisher Scientific) and mobile phase B was 0.1% formic acid (Thermo Fisher Scientific) in 80% acetonitrile/20% H_2_O (Thermo Fisher Scientific). Peptides passed through an Acclaim PepMap C18 100 Å, 3 μm, 75 μm × 2 cm trap column (Thermo Scientific) followed by separation on a PepSep C18 100 Å, 1.5 μm, 75 μm × 15 cm (Bruker) at a flow rate of 200 nl/min using the following 1 h gradient: 10%—35% B from 0 to 47 min, 35%—100% B from 47 to 55 min, 100% B from 55 min to 57 min, 100%–5% B from 57 min to 58 min, and 5% B from 58 min to 60 min. The trap column equilibration used a 9 μl at 3.0 μl/min flow rate and the separation column equilibration used a 12 μl at 3.0 μl/min flow rate. Additionally, 1 wash cycle of 20 μl and a flush volume of 100 μl were used. Peptides were ionized using the CaptiveSpray source, with a capillary voltage of 1650 V, dry gas flow of 3 l/min and temperature 180°C. Data were acquired using a positive ion mode diaPASEF method with a mass range from 100–1700 m/z and 1/K_0_ from 0.80 V_s_/cm^2^ to 1.35 V_s_/cm^2^ with 100 ms ramp time and 2 ms accumulation time. General tune parameters were: Funnel 1 RF = 300 Vpp, idCID Energy = 0 eV, Deflection Delta = 70 V, Funnel 2 RF = 200 Vpp, Multipole RF = 500 Vpp, Ion Energy = 5 eV, Low Mass = 200 m/z, Collision Energy = 10 eV, Collision RF = 1500 Vpp, Transfer Time = 60 μs, Pre Pulse Storage = 12 μs, and Stepping turned off. Tims tune parameters were: D1 = −20 V, D2 = −160 V, D3 = 110 V, D4 = 110 V, D5 = 0 V, D6 = 55 V, Funnel1 RF = 475 Vpp, and Collision Cell In = 300 V. Resulting spectra were uploaded to Spectronaut 18.7 (Biognosys, Cambridge, MA). Peptides were identified and quantified using the directDIA analysis default settings with the proteotypicity filter set to “Only Proteotypic” and a variable modification set to “methylthio.” MS1 protein group quantifications and associated protein group UniProt numbers and molecular weights were exported from Spectronaut (Biognosys). MS1 protein group quantifications, UniProt numbers, and molecular weights were imported into Perseus (Max Planck Institute of Biochemistry, Planegg-Martinsried, Bavaria, Germany) for use of the Proteomic Ruler plug-in, as previously described ^43^. The default proteomic ruler plug-in settings were used for hepatocytes with histone proteomic ruler as the scaling mode, ploidy set to two, and total cellular protein concentration set to 200 g/l. To account for differences in cell size, the total cellular protein concentration for endothelial cells, pericytes, and astrocytes was set to 100 g/l as a conservative estimate according to previous studies ^44^. The scaling mode and ploidy were set to the default proteomic ruler plug-in settings. Protein concentrations (nM) and copy numbers as estimated by the Proteomic Ruler were used for analysis.

#### *In Vitro* CKB-Mediated Tenofovir Enzyme activity assay

Primary human endothelial cells, pericytes, and astrocytes were lysed at 100% confluency with assay reaction buffer containing 75 mM HEPES pH 7.5 (Gibco), 5 mM MgCl_2_ (Thermo Fisher Scientific), 50 mM KCl (Sigma), 2 mM dithioerythritol (DTT) (Sigma), and LC-MS grade H_2_O (Thermo Fisher Scientific). Reactions were performed as previously described ^10^. In short, reactions were performed at a final volume of 200 μl in a reaction buffer containing: 75 mM HEPES pH 7.5, 50 mM KCl, 5 mM MgCl_2_, 2 mM DTT, 500 μg of total protein from lysate of brain endothelial cells, pericytes, or astrocytes, and 100 μM TFV-MP (Toronto Research Chemicals) to facilitate saturating conditions and to minimize the reverse reaction. Reaction solutions were prewarmed at 37°C for 5 minutes prior to initiation by the addition of initiation buffer containing 100 mM phosphocreatine or a combination of 100 mM phosphocreatine, 100 mM phosphoenolpyruvate, and 100 mM ATP for a final concentration of 500 μM phosphocreatine, phosphoenolpyruvate, and ATP. After 30 minutes of incubation at 37°C, reaction solutions were quenched with 200 μl of ice-cold 100% LC-MS methanol followed by brief vortex and incubation on ice for 10 minutes. Reactions were centrifuged at 10,000 rpm at 4°C for 10 minutes. Following centrifugation, the supernatants were removed and dried by vacuum centrifugation (Eppendorf, Hamburg, Germany). Control reactions were performed in parallel by omitting the addition of initiation buffer, cell lysate or TFV-MP. Independent experimental assay reactions were completed for endothelial cells (n=3-5), pericytes (n=4), and astrocytes (n=4-5).

TFV-DP formation was measured by triple quadrupole LC-MS/MS. Dried samples were reconstituted in 30 μl of 0.1% formic acid in water (v/v) (Thermo Fisher Scientific). Samples were subsequently analyzed on an Agilent Ultivo Triple Quadrupole LC/MS (Agilent, Santa Clara, CA). Mobile phase A was 5 mM *N*,*N*-dimethylhexylamine (DMHA) (Sigma) in water at pH 7.0, and mobile phase B was 5 mM DMHA in 50% acetonitrile/50% water (v/v). Analytes were separated on a Halo C18 column (2.1 mm x 100 mm, 2.1 µm, Mac-Mod Analytical) (Mac-Mod Analytical, Chadds Ford, PA) at a flow rate of 0.450 mL/min using the following gradient: 5% B from 0 to 1 minutes, 5%–45% B from 1 to 9 minutes, 45% B from 9 to 13 minutes, 45%–100% B from 13 to 15 minutes, 100% B from 15 to 20 minutes, 100%–5% B from 20 to 23 minutes, and 5% B from 23 to 30 minutes. Analytes were ionized by heated electrospray ionization with a capillary voltage of 4800 V, gas temperature of 260°C, gas flow rate of 11.0 L/min, and nebulizer pressure of 30 psi. TFV was detected in positive mode using a single-reaction monitoring scan with a transition of mass to charge ratios of 288.1 **→** 176 with a scan/dwell time of 400 ms, fragment energy of 186 V, and collision energy of 27 V. TFV-MP was detected in positive mode using a single-reaction monitoring scan with a transition of mass to charge ratios of 368.1 **→** 270 with a scan/dwell time of 400 ms, fragment energy of 181 V, and collision energy of 19 V. TFV-DP was detected in positive ion mode using a single-reaction monitoring scan with a transition of mass to charge ratios of 448 → 350 with a scan/dwell time of 400 ms, fragment energy of 230 V, and collision energy of 15 V. Chromatographic peak areas were used for relative comparisons of metabolite abundance.

#### Efflux transporter activity assay and flow cytometry

Efflux transporter activity was determined by measuring intracellular fluorescence following uptake of rhodamine 123 (10 µM, Thermo Fisher Scientific), Hoechst 33342 (5 µg/mL, Thermo Fisher Scientific), and monobromobimane (10 µM, Thermo Fisher Scientific), fluorescent substrates with specificity for P-gp, BCRP, and MRP4, respectively. Hoechst 33342 is a well-established transport substrate of BCRP; however, evidence exists that there is substrate specificity overlap with P-gp, which may contribute to mixed efflux effects ^49^. We did not observe intracellular trapping of monobromobimane or oxidative stress to induce cell death (data not shown). Primary human brain endothelial cells, pericytes, and astrocytes were seeded onto 6-well plates at a seeding density of 0.3 x 10^6^ cells/well. Upon reaching 100% confluency, cells were incubated with each fluorescent substrate for 15 minutes at 37°C, 5% CO_2_ to allow uptake into the cell, after which fresh media was added, and the fluorescent substrates allowed to efflux from the cells for two hours at 37°C, 5% CO_2_. Addition of fluorescent substrate for 15 minutes at 37°C, 5% CO_2_ that was not permitted to efflux was used as a positive control to determine maximal substrate uptake. Following exposure to the fluorescent substrates, cells were washed with PBS (Gibco), filtered using BD FACS tubes (BD Biosciences, Franklin Lakes, NJ) with cell strainer caps with 35-µm pores (BD Biosciences), and immediately subject to flow cytometric acquisition where at least 10,000 singlet events were acquired with a BD LSRFortessa cytometer and Diva software version 9 on the Windows 10 platform (BD Biosciences). Flow cytometric data were analyzed using FlowJo version 10.9 (FlowJo, Ashland, OR) where fluorescence intensity of substrates in cells that received substrate for two hours (efflux) was subtracted from the cells that received substrate at the end of the two hours (uptake) to quantitate efflux capacity.

#### Cell viability determination

Brain endothelial cells, pericytes, and astrocytes were exposed to rhodamine 123 (10 µM, Thermo Fisher Scientific), Hoechst 33342 (5 µg/mL, Thermo Fisher Scientific), and monobromobimane (10 µM, Thermo Fisher Scientific) for 15 minutes and two hours, after which time viability was assessed, as previously described ^18^, using the BD Horizon Fixable Viability Stain 520 (BD Biosciences), according to manufacturer’s instructions. Heat shock at 55°C for 10 minutes served as a positive control for cell death. The cells were analyzed by flow cytometry within 5 minutes of staining.

#### Thymidine kinase activity assay

To determine thymidine kinase activity, the DiviTum^TM^ (Biovica International, San Diego, CA) thymidine kinase activity assay was performed according to manufacturer’s instructions, as previously described ^50^. When 100% confluent, endothelial cell, pericyte, and astrocyte monoculture cells were lysed with DiviTum^TM^ TK activity lysis buffer (Biovica International). Cell lysates were mixed with reaction mixture in an enzyme-linked immunosorbent assay (ELISA) titer plate where bromodeoxyuridine (BrdU) monophosphate is generated by the TK reaction, which is phosphorylated to BrdU triphosphate. BrdU triphosphate was detected by ELISA using an anti-BrdU monoclonal antibody that is conjugated to alkaline phosphatase and a chromogenic substrate, producing the optical density of the color. The absorbance readings to DiviTum units per liter (Du/L) were converted using the values from standards with known TK activity, with a working range from 20 to 4000 Du/L. TK activity was reported as DiviTum TK activity value (DuA) per total protein (µg). The assays were performed at the Biovica laboratory in San Diego, California, and investigators were blinded to cell-type. Thymidine kinase activity was normalized to individual protein amount in brain endothelial cell, pericyte, and astrocyte lysates as measured by the Pierce^TM^ Bicinchoninic acid (BCA) assay (Thermo Fisher Scientific).

#### Pyruvate kinase activity assay

Pyruvate kinase activity was determined using the pyruvate kinase activity assay kit (Abcam, Cambridge, MA) according to the manufacturer’s instructions. When 100% confluent, primary human brain endothelial cells, pericytes, and astrocytes were lysed using cold pyruvate kinase assay buffer. Lysates were diluted using pyruvate kinase assay buffer and a colorimetric assay performed where the absorbance of the oxidized pyruvate was measured at 570 nm at 25°C. Absorbance readings were taken by Synergy HT microplate reader (BioTek, Winooski, VT) every minute for 20 minutes. Pyruvate kinase activity was calculated using the concentration of pyruvate from the pyruvate standard curve generated in the assay over 20 minutes and the volume of lysate added to the 96-well for the assay. The sensitivity of this assay is 0.1 mU/mL. Pyruvate kinase activity was normalized to individual protein amount in brain endothelial cell, pericyte, and astrocyte lysates as measured by the Pierce^TM^ Bicinchoninic acid (BCA) assay (Thermo Fisher Scientific).

#### Blood brain barrier cell HIV and ART exposure

Primary human brain endothelial cells and astrocytes were seeded on 10 cm plates at a density of 1 x 10^6^ cells/plate. Primary human pericytes were seeded on 150 cm^2^ flask at a density of 1 x 10^6^ cells/flask. Upon reaching 90% confluency, cells were washed with PBS (Gibco) and treated with ART, including TFV (Toronto Research Chemicals), FTC (Toronto Research Chemicals), and DTG (Toronto Research Chemicals), HIV_ADA_ (5 ng/mL) (National Institutes of Health), ART in combination with HIV, or vehicle for 24 hours at 37°C, 5% CO_2_. All treatments were constituted in cell culture media to reach final treatment concentrations. Final ART concentrations were 10 µM, which reflects the maximum serum concentrations (C_max_) of drug obtained from clinical studies ^15^. Following treatment, cells were washed twice with PBS (Gibco), centrifuged, and lysed in 1X proteomics lysis buffer, containing 5% sodium dodecyl sulfate (SDS) (Invitrogen) and 50 mM triethylammonium bicarbonate (Sigma). Cell lysates were processed for proteomics as described above.

#### SimpliFi Pathway Analysis of HIV/ART-exposed BBB cells

Spectronaut (Biognosys) proteomics analysis, including MS1 protein group quantities, of vehicle, HIV, ART, and HIV+ART-exposed brain endothelial cells, pericytes, and astrocytes was imported into the internet-based platform SimpliFi (ProtiFi) and assigned into experimental batches. SimpliFi quality control parameters found no batch effects between experimental batches. Minimal run order effects were detected between batches. Pathway hits for each cell type and exposure condition were filtered by p-value less than 0.05 and log2 fold change greater than 1, and pathway hits of interest were subsequently selected using these parameters.

#### Blood brain barrier proteomics repository

The Human Blood Brain Barrier Monoculture Proteome Repository 1.0.2 was developed with base R (R version 4.2.2 (2022-10-31)) in the MacOS operating environment with visualization provided through the Shiny application framework and the Shiny Dashboard R package. The dependencies and their respective versions are as follows: “scales_1.2.1”, “shinydashboard_0.7.2”, “shinythemes_1.2.0”, “readxl_1.4.2”, “janitor_2.2.0”, “DT_0.28”, “tidyr_1.3.0”, “dplyr_1.1.3”, “readr_2.1.4”, “ggpubr_0.6.0”, “readxl_1.4.2”, “plotly_4.10.2”, “ggplot2_3.4.3”, “ggpattern_1.0.1”, and “shiny_1.7.4.1”. The repository includes data summarized in Figure 2. Brain endothelial cell, pericyte, astrocyte, and hepatocyte proteomics were processed by Spectronaut (Biognosys) and using the Proteomic Ruler plug-in in Perseus (Max Planck Institute of Biochemistry) as described above ^43^. The missing values, NA and NaN, were replaced with zeroes. The Human Blood Brain Barrier Monoculture Proteome Repository 1.0.2 was made publicly available:(https://hannahwilkins.shinyapps.io/Blood_Brain_Barrier_Cells_Proteomics_Repository_Publish/).

#### Quantification and Statistical analysis

At least three independent experiments were performed for all *in vitro* experiments. Frontal cortex, thalamus, and cerebellum tissues were collected from two rhesus macaques due to sample availability. Rhesus macaque astrocytes were collected from five independent rhesus macaque brains. Liquid chromatography/mass spectrometry and proteomic assays were run with three independent sample injections. Details regarding the number of independent experiments performed are included in all figure legends. All statistical analyses were presented as mean (±SD) for at least three independent experiments, with the exception of rhesus macaque brain region ART analyses (Figure 1A-B) which were limited to two independent rhesus macaques due to sample availability. Statistical analyses were performed using prism software 10.1.1 GraphPad Software, Inc., San Diego, CA. A D’Agostino-Pearson normality test was performed as appropriate to evaluate whether the data fit a Gaussian distribution. When the data were normally distributed, a Brown-Forsythe and Welch ANOVA with a Dunnett’s T3 multiple comparisons test was performed as appropriate. When the data were not normally distributed, a Kruskal-Wallis test with Dunn’s multiple comparisons test was performed as appropriate. For the efflux transporter activity assay, a paired t-test was performed between efflux and uptake, and for BBB cell comparisons in efflux capacity, an ordinary one-way ANOVA was performed. For SimpliFi (ProtiFi) pathway analysis of vehicle, HIV, ART, and HIV+ART-exposed brain endothelial cells, pericytes, and astrocytes, statistics were performed by the internet-based SimpliFi (ProtiFi) platform. For proteomics by LC-MS/MS data, values reported as “NaN” by Spectronaut (Biognosys) or SimpliFi (ProtiFi) were reported as missing and recorded as 0. Experimental or technical replicates with missing LC-MS/MS values were omitted from statistical analyses. >2 independent experiments with missing values were reported as not reliably detected (ND) by proteomics analyses. When present, the vehicle treatment condition served as the reference group for multiple comparison analyses in the one-way ANOVA test. *p≦0.05. **p≦0.01. ***p≦0.001 ****p≦0.0001.

