## Supplemental Figures S1-S6 for "Drug Metabolism and Transport Capacity of Endothelial Cells, Pericytes, and Astrocytes: Implications for CNS Drug Disposition"

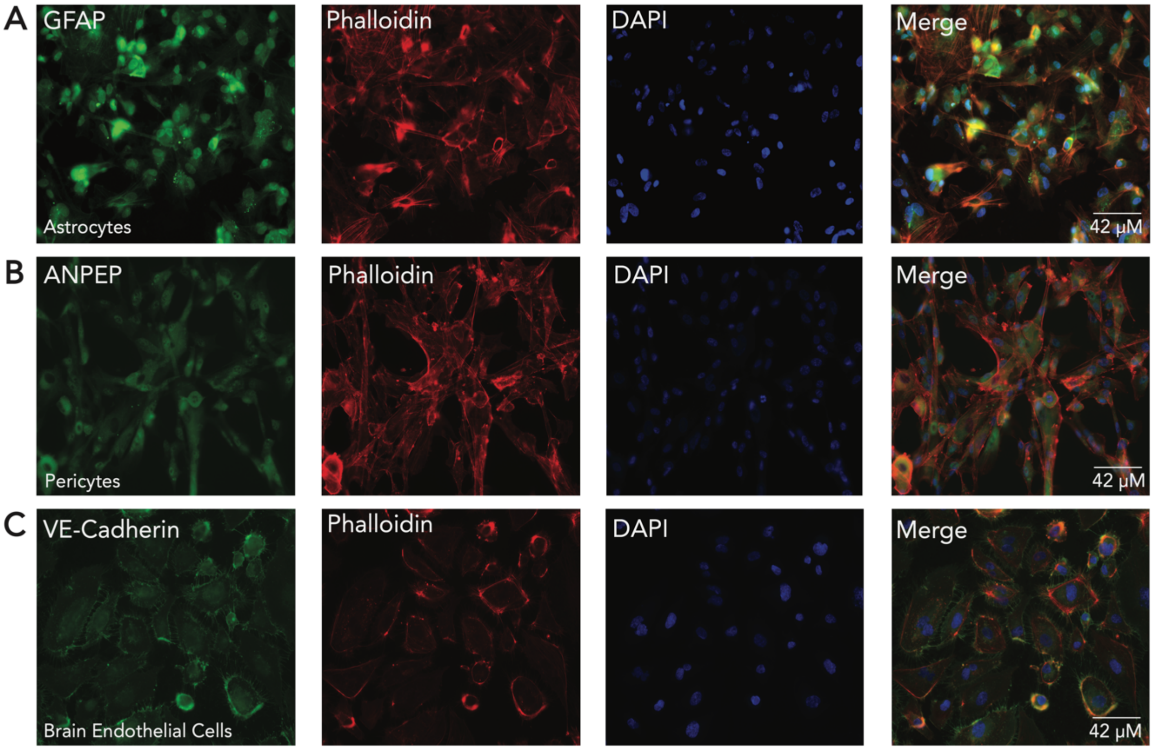


**Figure S1. Brain Microvascular Endothelial Cells (BMVEC), pericytes and astrocytes express characteristic cell markers.** Immunofluorescent microscopy was performed to evaluate expression of anticipated markers in primary human **(A)** astrocytes, **(B)** pericytes, and **(C)** BMVEC. Antibodies with specificity to **(A)** GFAP, **(B)** ANPEP, and **(C)** VE-Cadherin were coupled to Alexa Fluor 488 for analysis. DAPI was used to visualize the nucleus, and phalloidin was used to visualize actin filaments. Merge depicts the combined signal for proteins of interest (green), phalloidin (red), and DAPI (blue). Representative images, out of 20 independent images, are shown. All scale bars = 42 μm, and all images are at the same scale.

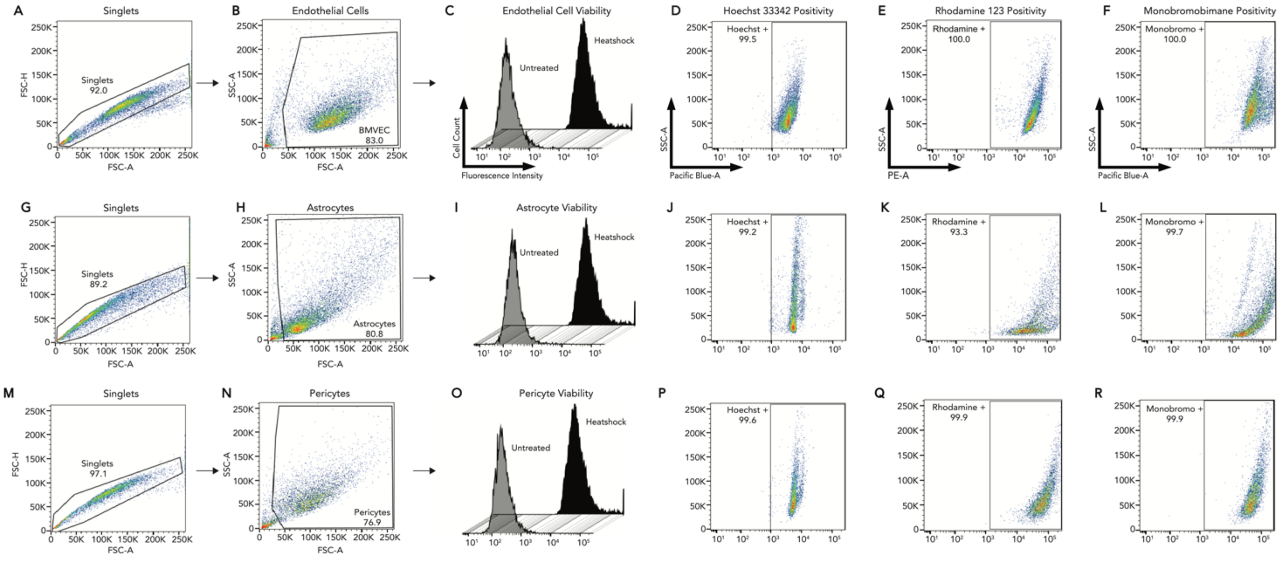


**Figure S2. Representative flow cytometric analyses of primary human blood brain barrier cell-based efflux transporter activity assay.** Doublets were excluded using FSC-H and FSC-A measurements for **(A)** brain endothelial cells, **(G)** astrocytes, and **(M)** pericytes. Debris was gated out by drawing a gate on cell-sized events using SSC-A and FSC-A measurements for **(B)** endothelial cells, **(H)** astrocytes, and **(N)** pericytes. **(A)** Endothelial cell, **(I)** astrocyte, and **(O)** pericyte viability was assessed using heat shock at 56°C for 30 minutes as a positive control (black histogram) compared to untreated, live cells (gray histogram). Hoechst 33342 positive cells were gated using SSC-A and Pacific Blue-A measurements for **(D)** brain endothelial cells, **(J)** astrocytes, and **(P)** pericytes. Hoechst 33342 positive cell populations were expressed as a percentage. Rhodamine 123 positive cells were gated using SSC-A and PE-A measurements for **(E)** endothelial cells, **(K)** astrocytes, and **(Q)** pericytes. Rhodamine 123 positive cell populations were expressed as a percentage. Monobromobimane positive cells were gated using SSC-A and Pacific Blue-A measurements for **(F)** endothelial cells, **(L)** astrocytes, and **(R)** pericytes. Monobromobimane positive cell populations were expressed as a percentage.

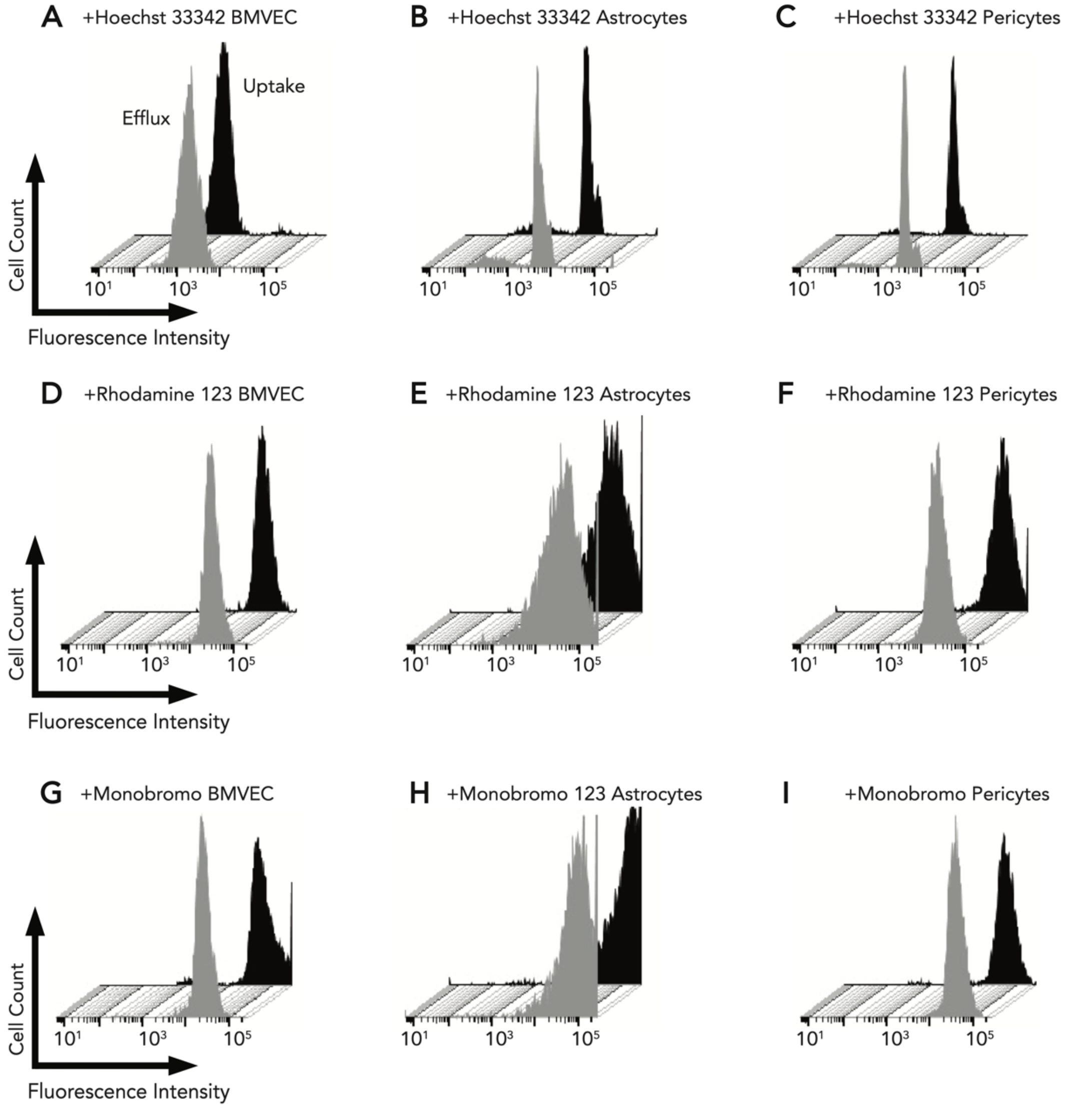


**Figure S3. Representative histograms of efflux and uptake of substrate in primary human blood brain barrier cell-based efflux transporter activity assays.** Efflux (gray histogram) and uptake (black histogram) of Hoechst 33342 in **(A)** endothelial cells (BMVEC), **(B)** astrocytes, and **(C)** pericytes. Efflux (gray histogram) and uptake (black histogram) of Rhodamine 123 in **(D)** endothelial cells (BMVEC), **(E)** astrocytes, and **(F)** pericytes. Efflux (gray histogram) and uptake (black histogram) of monobromobimane in

**(G)** endothelial cells (BMVEC), **(H)** astrocytes, and **(I)** pericytes. For all cells, uptake was measured after 15 minutes of incubation with substrate at 37 °C, 5% $CO_2$, and efflux was measured over 2 hours at 37 °C, 5% $CO_2$.

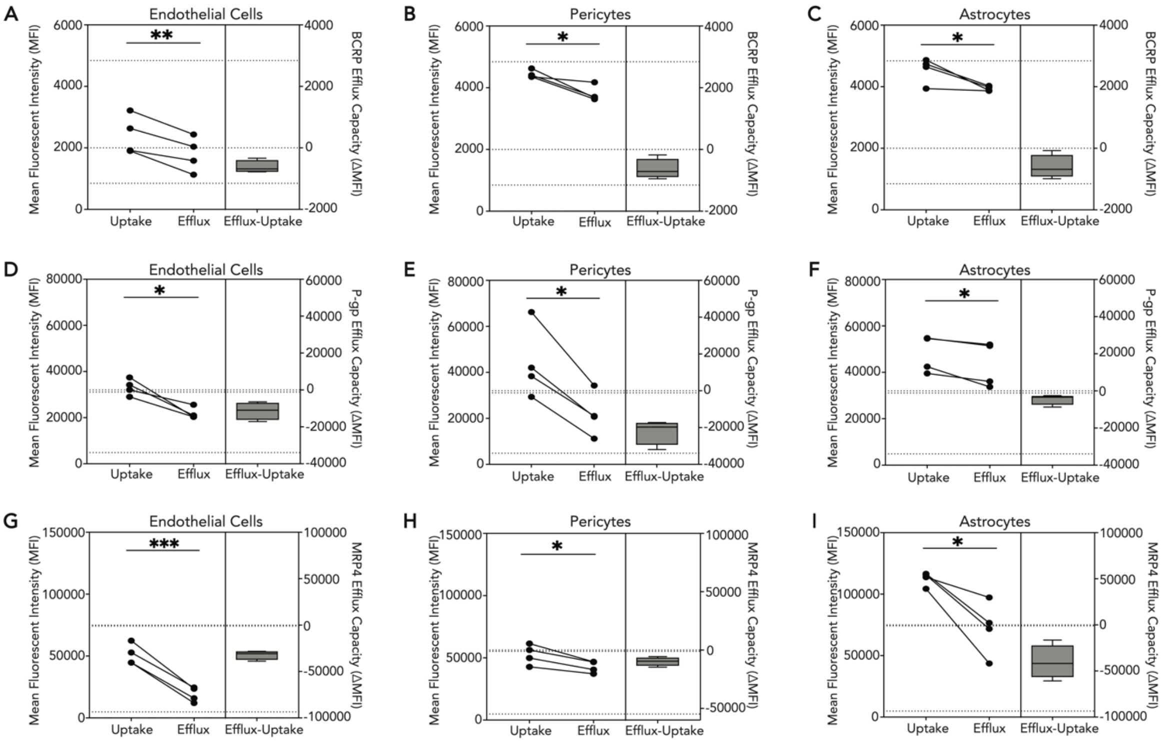


**Figure S4. TFV/FTC efflux transporters are differentially active in blood brain barrier (BBB) cells.** Transporter efflux was measured in primary human BBB cells where monocultures of endothelial cells (BMVEC), pericytes, and astrocytes were loaded in select wells of a 6-well plate with dyes specific for **(A-C)** BCRP (Hoechst 33342, 5 µg/mL), **(D-F)** P-gp (rhodamine 123, 10 µM), and **(G-I)** MRP4 (monobromobimane, 10 µM) for 15 minutes at 37 °C, 5% $CO_2$. The cells were washed by replacing the media, and the dyes were subsequently allowed to efflux out of the cell for 2 hours at 37 °C, 5% $CO_2$. Uptake was measured by loading select wells of a 6-well plate with Hoechst 33342, rhodamine 123, or monobromobimane for 15 minutes 37 °C, 5% $CO_2$. Flow cytometric analysis was performed to evaluate the fluorescence of substrates. Mean fluorescence intensities of substrate uptake and efflux were summarized by the difference in uptake and efflux (efflux capacity) for BCRP in **(A)** endothelial cells, **(B)** pericytes, and **(C)** astrocytes, for P-gp in **(D)** endothelial cells, **(E)** pericytes, and **(F)** astrocytes as well as for MRP4 in **(G)** endothelial cells, **(H)** pericytes, and **(I)** astrocytes. Four independent experiments were performed, and statistical analysis was conducted by paired t test. *$p < 0.05$, **$p < 0.01$, ***$p<0.001$.

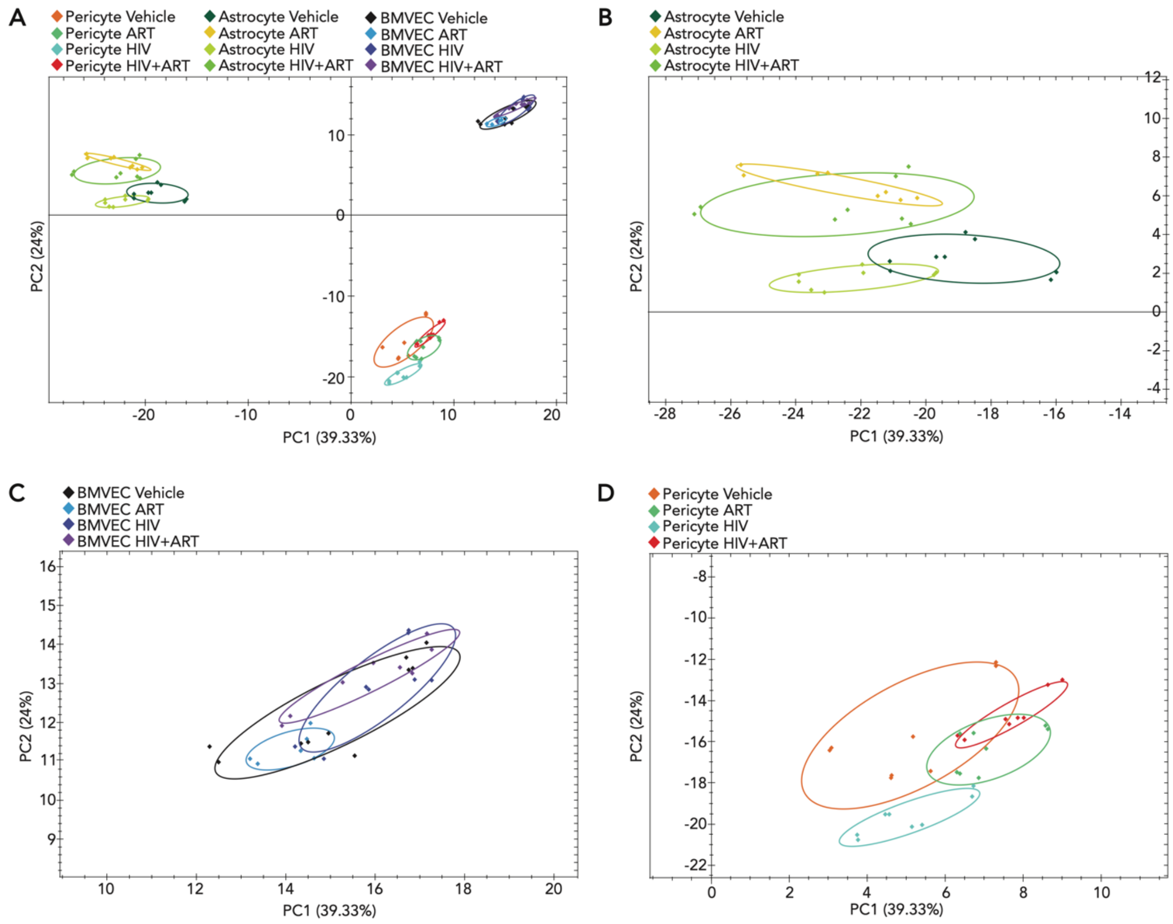


**Figure S5. Principal Component Analysis (PCA) of brain microvascular endothelial cells (BMVEC), pericytes, and astrocytes exposed to ART, HIV, and a combination of HIV and ART (HIV+ART).** Primary human pericyte, BMVEC, and astrocyte monocultures were exposed to ART (10 µM TFV, 10 µM FTC, 10 µM DTG), HIV (5 ng/mL), or a combination of HIV and ART (10 µM TFV, 10 µM FTC, 10 µM DTG, and 5 ng/mL HIV) for 24 h at 37 °C, 5% $CO_2$. Exposure to vehicle was used as a control. Pericytes, BMVEC, and astrocytes were lysed and processed for proteomics analyses. **(A)** PCA plot was generated by Spectronaut proteomics processing software to indicate sample clustering by exposure condition (Vehicle, HIV, ART, and HIV+ART) for pericytes, BMVEC, and astrocytes. PCAs were zoomed in to specific cell type within

**(A)** to visualize individual cell type clustering for **(B)** astrocytes, **(C)** BMVEC, and **(D)** pericytes.

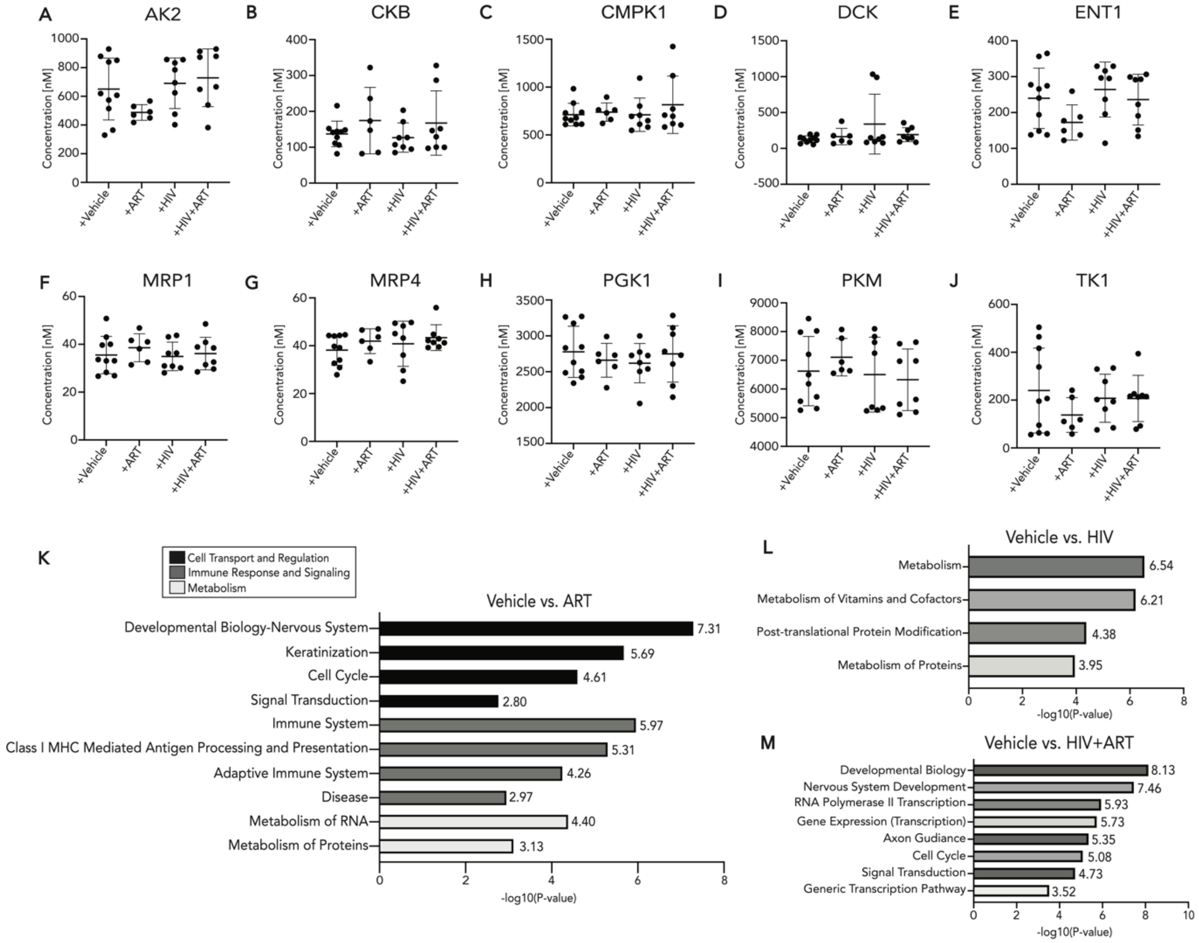


**Figure S6. Endothelial cell proteomes and cellular pathways are minimally impacted by ART, HIV, and a combination of HIV and ART**. Primary human endothelial cell (BMVEC) monocultures were exposed to ART (10 µM TFV, 10 µM FTC, 10 µM DTG), HIV (5 ng/mL), or HIV+ART (10 µM TFV, 10 µM FTC, 10 µM DTG, 5 ng/mL HIV) for 24 h at 37 °C, 5% $CO_2$. Treatment with vehicle was used as a control. BMVEC were lysed and processed for proteomics analyses. Concentrations of proteins are measured by Proteomic Ruler. TFV/FTC transporter and nucleotide metabolizing kinase protein concentrations were measured for **(A)** AK2, **(B)** CKB, **(C)** CMPK1, **(D)** DCK, **(E)** ENT1, **(F)** MRP1, **(G)** MRP4, **(H)** PGK1, **(I)** PKM, and **(J)** TK1. Three to five independent experiments with two LC-MS/MS injection replicates each were performed per condition (represented by individual dots). Replicates with missing LC-MS/MS values were omitted from plot. Data represented as mean ± standard deviation. Statistical analysis was performed by a Brown-Forsythe and Welch ANOVA or a Kruskal-Wallis ANOVA. Pathways that were significantly altered after **(K)** ART, **(L)** HIV, or **(M)** HIV+ART exposure (compared to vehicle) in BMVEC were reported by SimpliFi proteomics software. **(K)** Pathways of interest were only identified for BMVEC exposed to ART. All pathway hits were displayed for BMVEC exposed to **(L)** HIV and **(M)** HIV+ART. Pathways of interest were filtered by pathway changes greater than one log fold and hypergeometric p-value $< 0.05$. Pathways were ranked by –log10(hypergeometric p-value). Pathway analyses summarize three to five independent experiments per condition with two LC-MS/MS injection replicates each.
