## Supplementary material for "Drug Metabolism and Transport Capacity of Endothelial Cells, Pericytes, and Astrocytes: Implications for CNS Drug Disposition": Key Resources Table

| Reagent or Resource | Source | Identifier |
|---|---|---|
| Antibodies | | |
| ABCG2/CD338 BCRP;<br>Clone 3G8 | NovusBio | Cat#: NBP2-22124 |
| P-gp polyclonal antibody | Invitrogen | Cat#: PA5-61300<br>RRID: AB_2645457 |
| MRP4/ABCC4 Rabbit mAb | Cell Signaling Technology | Cat#: 12705 |
| Anti-MRP1 antibody<br>Clone MRPm5 | Abcam | Cat#: Ab24102 |
| ENT1 polyclonal antibody | Invitrogen | Cat#: PA5-116461<br>RRID: AB_2901093 |
| Beta-Actin Mouse mAb<br>(HRP Conjugate); IgG2b | Cell Signaling Technology | Cat#: 8H10D10 |
| PGK1 PolyClonal Antibody | Invitrogen | Cat#: PA5-28612<br>RRID: AB_2546088 |
| AK2 Polyclonal Antibody | Proteintech | Cat#: 11014-1-AP<br>RRID: AB_2305358 |
| CKB/CKM Polyclonal Antibody | Proteintech | Cat#: 15137-1-AP<br>RRID: AB_2080878 |
| Anti-creatine kinase B type antibody (Knockout validated) | Abcam | Cat#: Ab126418 |
| TK1 Polyclonal antibody | Proteintech | Cat#: 15691-1-AP<br>RRID: AB_2203613 |
| UMP/CMP kinase polyclonal antibody | Proteintech | Cat#: 11360-1-AP<br>AB_2081395 |
| DCK Polyclonal Antibody | Proteintech | Cat#: 17758-1-AP<br>RRID: AB_2261476 |
| PKM1-specific Polyclonal antibody | Proteintech | Cat#: 15821-1-AP<br>RRID: AB_2163820 |
| PKLR Polyclonal antibody | Proteintech | 2 Cat#: 2456-1-AP<br>RRID: AB_10918271 |
| Goat Anti-Mouse IgG H&L (HRP) secondary antibody | Abcam | Cat#: ab97023 |
| Goat Anti-Rabbit IgG H&L (HRP) | Abcam | Cat#: Ab97051 |
| GFAP (AF488) | Invitrogen | Cat#: 53-9892-82<br>RRID: AB_10598515 |
| CD13 (ANPEP) Recombinant Rabbit Monoclonal Antibody (SC70-01) | Invitrogen | Cat#: MA5-32226<br>RRID: AB_2809512 |

| | | |
|---|---|---|
| CD144 (VE-cadherin) Monoclonal Antibody (16B1) | Invitrogen | Cat#: 14-1449-82<br>RRID: AB_467495 |
| Goat anti-Rabbit IgG (H+L) Cross-Absorbed Secondary Antibody, Alexa Fluor™ 488 | Abcam | Cat#: ab97051<br>RRID: AB_143165 |
| Goat anti-mouse IgG (H+L) Cross-Absorbed Secondary Antibody, Alexa Fluor™ 488 | Abcam | Catalog #: A-11001<br>RRID: AB_2534069 |
| Chemicals and Reagents | | |
| Basal Medium Eagle (BME) | Thermo Fisher Scientific: Gibco | Cat#: 21010046 |
| Sodium bicarbonate | Sigma-Aldrich | Cat#: S5761 |
| Poly-L-lysine solution | Sigma-Aldrich | Cat#: P8920 |
| HEPES (1 M) | Thermo Fisher Scientific: Gibco | Cat#: 15630080 |
| Fetal Bovine Serum | R&D Systems | Cat#: S11150 |
| Penicillin-Streptomycin (10,000 U/mL) | Thermo Fisher Scientific: Gibco | Cat#: 15140122 |
| Astrocyte Growth Supplement | ScienCell | Cat#: 1852 |
| Gelatin (10% soln) | Thermo Scientific Chemicals | Cat#: J62699.AK |
| Medium 199 | Thermo Fisher Scientific: Gibco | Cat#: 11150067 |
| Newborn Calf Serum, heat inactivated, New Zealand origin | Thermo Fisher Scientific: Gibco | Cat#: 26010074 |
| Heparin sodium | Milipore Sigma | Cat#: 9041-08-1 |
| GemCell™ U.S. Origin Human Serum AB | GeminiBio | Cat#: 100-512-100 |
| L-Ascorbic acid | Milipore Sigma | Cat#: A92902 |
| Endothelial Cell Growth Supplement | Milipore Sigma | Cat#: 02-102 |
| L-Glutamine | Thermo Fisher Scientific: Gibco | Cat#: 25030081 |
| Bovine Brain Extract | Lonza | Cat#: CC-40908 |
| Pericyte Medium | ScienCell | Cat#: 1201 |
| Pericyte Growth Supplement | ScienCell | Cat#: 1252 |
| Hanks' Balanced Salt Solution (HBSS) without Ca2+, Mg2+ for Pierce™ Primary Cell Isolation Kits | Thermo Fisher Scientific | Cat#: 88284 |
| DMEM, high glucose | Thermo Fisher Scientific: Gibco | Cat#: 11965092 |
| Geneticin Selective Antibiotic (G418 Sulfate) (50 mg/mL) | Thermo Fisher Scientific: Gibco | Cat#: 10131027 |

| | | |
|---|---|---|
| Sodium Pyruvate (100 mM) | Thermo Fisher Scientific: Gibco | Cat#: 11360070 |
| PBS, pH 7.4 | Thermo Fisher Scientific: Gibco | Cat#: 10010023 |
| DPBS, no calcium, no magnesium | Thermo Fisher Scientific: Gibco | Cat#: 14190144 |
| DNase I | Invitrogen | Cat#: 18047019 |
| Trypsin (2.5%), no phenol red | Thermo Fisher Scientific: Gibco | Cat#: 15090046 |
| Cytiva Percoll™ Centrifugation media | Fisher Scientific | Cat#: 45-001-748 |
| HEPES, 1.0M buffer solution, pH 7.5 | Thermo Scientific Chemicals | Cat#: J60712.AP |
| Adenosine 5'-triphosphate disodium salt trihydrate | Milipore Sigma | Cat#: 10127523001 |
| Potassium Chloride | Sigma-Aldrich | Cat#: P3911 |
| Water, Optima™ LC/MS Grade, Fisher Chemical™ | Fisher Scientific | Cat#: W6500 |
| Methanol, Optima™ LC/MS Grade, Fisher Chemical™ | Fisher Scientific | Cat#: A456-500 |
| Water with 0.1% Formic acid (v/v), Optima™ LC/MS Grade, Thermo Scientific™ | Fisher Scientific | Cat#: LS118-500 |
| Acetonitrile, Optima™ LC/MS Grade, Fisher Chemical™ | Fisher Scientific | Cat#: A955-500 |
| N,N-Dimethylhexylamine (DMHA) | Sigma-Aldrich | Cat#: 308102 |
| Trifluroacetic acid | Sigma-Aldrich | Cat#: T6508 |
| Dithioerythritol (DTT) | Milipore Sigma | Cat#: 10197777001 |
| Creatine phosphate, disodium salt | Fisher Scientific | Cat#: ICN10052001 |
| Phosphoenolpyruvic acid monopotassium slat | Fisher Scientific | Cat#: AAB2035806 |
| Tenofovir | Toronto Research Chemicals | Cat#: T018500 |
| Tenofovir Monophosphate | Toronto Research Chemicals | Cat#: T018560 |
| Tenofovir Diphosphate | Toronto Research Chemicals | Cat#: T018525 |
| Emtricitabine | LGC Standards | Cat#: TRC-E525000 |
| Dolutegravir | LGC Standards | Cat#: TRC-D528800 |
| RIPA Buffer (10X) | Cell Signaling Technology | Cat#: 9806 |
| Protease/Phosphatase Inhibitor Cocktail (100X) | Cell Signaling Technology | Cat#: 5872 |

| | | |
|---|---|---|
| Bio-Rad Protein Assay Dye Reagent Concentrate | Bio-Rad | Cat#: **5000006** |
| Dry Milk, Non-Fat, Molecular biology Grade | Lab Scientific | Cat#: MSPP-M0841 |
| Tris Buffered Saline (TBS) (10X), pH 7.4 | Quality Biological | Cat#: 351-086-101 |
| Tween-20 | Sigma-Aldrich | Cat#: P9416 |
| Revvity Health Sciences Inc Western Lightning Plus-ECL, Enhanced Chemiluminescence Substrate | PerkinElmer | Cat#: 50-904-9326 |
| Bovine Serum Albumin (BSA) | Thermo Fisher Scientific | Cat#: B14 |
| Restore™ PLUS Western Blot Stripping Buffer | Thermo Fisher Scientific | Cat#: 46430 |
| Triethylammonium bicarbonate buffer | Sigma-Aldrich | Cat#: T7408 |
| UltraPure™ SDS Solution, 10% | Invitrogen | Cat#: 15553027 |
| Phosphoric Acid | Sigma-Aldrich | Cat#: 695017 |
| Magnesium Chloride, anhydrous, 99% | Thermo Fisher Scientific | Cat#: 012315.C4 |
| Pierce Trypsin Protease, MS Grade | Thermo Fisher Scientific | Cat#: 90057 |
| TrypLE™ Express Enzyme (1X), no phenol red | Thermo Fisher Scientific: Gibco | Cat#: 12604103 |
| Paraformaldehyde 4% Aqueous Solution, EM Grade | Electron Microscopy Sciences | Cat#: 157-4-1L |
| Wheat Germ Agglutinin (WGA) | Invitrogen | Cat#: W21405 |
| Triton™ X-100 | Sigma-Aldrich | X Cat#: 100-5ML |
| Ethylenediaminetetraacetic acid, (EDTA) | Milipore Sigma | Cat#: E9884 |
| Gelatin from cold water fish skin | Milipore Sigma | Cat#: 9000-70-8 |
| Bovine Serum Albumin | Sigma-Aldrich | Cat#: A2058 |
| Normal Goat Serum Blocking Solution | Vector Laboratories | Cat#: S-1000-20 |
| Ibidi Mounting Medium with DAPI | Ibidi | Cat#: 50011 |
| Trypsin-EDTA (0.5%), no phenol red | Thermo Fisher Scientific | Cat#: 15400054 |

| | | |
|---|---|---|
| Rhodamine 123, 25 mg | Invitrogen | Cat#: R302 |
| Hoechst 33342 Ready Flow™ Reagent | Thermo Fisher Scientific | Cat#: R37165 |
| Monobromobimane (mBBr), FluoroPure™ grade | Invitrogen | Cat#: M20381 |
| BD Horizon™ Fixable Viability Stain 520 | BD Bioseicnes | Cat#: 564407 |
| Critical Commercial Assays | | |
| Universal proteomics sample preparation kits, minis (100 100µg - 300 µg) with solutions | Protifi | Cat#: K02-mini-10 |
| Pierce™ Bradford Protein Assay Kit | Thermo Scientific | Cat#: 23200 |
| Pierce™ BCA Protein Assay Kits | Thermo Scientific | Cat#: 23225 |
| Pyruvate Kinase Assay Kit | Abcam | Cat#: aPb83432 |
| Pierce™ Quantitative Peptide Assays and Standards | Thermo Scientific | Cat#: 23275 |
| Thymidine Kinase assay | DiviTum TKa<br>Rae et al., 2023 | NA |
| HIV p24 ELISA | Perkin Elmer | Cat#: NEK050 |
| Experimental Models: Cell Lines | | |
| Primary human astrocytes | ScienCell | Cat#: 1800 |
| Primary human vascular pericytes | ScienCell | Cat#: 1200 |
| Primary human brain microvascular endothelial cells | Cell Systems | Cat#: ACBRI 376 |
| Fresh primary human hepatocyte suspension | BioIVT | Cat#: F00986 |
| Virus Strains | | |
| SIVmac251 | Abreu et al., 2019 | N/A |
| Human Immunodeficiency Virus-1 (HIV-1) ADA | NIH HIV Reagent Program/BEI Resources Repository, Division of AIDS, NIAID, NIH | ARP-416 |
| Experimental Models | | |
| Rhesus macaques | The Johns Hopkins University School of Medicine rhesus colony | N/A |
| Deposited Data | | |

| | | |
|---|---|---|
| Software and algorithms | | |
| GraphPad Prism v10 | GraphPad | N/A |
| Perseus v2.0.7.0; Proteomic Ruler Plug-in | MaxQuant; Max-Planck Institute of Biochemistry, Computational Systems Biochemistry citation | N/A |
| SimpliFi | Pappin et al., 2020 | N/A |
| Spectronaut v18.7 | Biognosys | N/A |
| Flowjo v10 | Flowjo | N/A |
| R Studio | 2023.12.1 | N/A |
| Fiji v2.14.0 | Fiji | N/A |
| Other | | |
| Cell Culture Flask, T-150 | StemCell Technologies | Cat#: 38072 |
| 10 cm Culture Dish | StemCell Technologies | Cat#: 38069 |
| Falcon ® 6-Well Flat-Bottom Plate, Tissue Culture-Treated | StemCell Technologies | Cat#: 38016 |
| Acclaim$^{TM}$ PepMap$^{TM}$ 100 C18 HPLC Columns | Thermo Scientific | Cat#: 164946 |
| PepSep C18 100 Å, 1.5 µm, 75 µm × 15 cm | Bruker Daltonics | Cat#: 1893625 |
| HALO C18, 90 Å. 2.7 µm, 2.1 x 100 mm Fused-Core Column | MacMod Analytical | Cat#: 92812602 |
| Corning ® Cell Strainer, 100 µm Nylon | Corning | Cat#: 431752 |
| Falcon Cell Strainer 40 µm Nylon | Corning | Cat#: 352340 |
| Falcon Cell Strainer 70 µm Nylon | Corning | Cat#: 352350 |
| Cell Culture Imaging Dish µ-Dish 35 mm, high, ibiTreat | Ibidi | Cat#: 81156 |

| | | |
|---|---|---|
| 4–12% Criterion™ XT Bis-Tris Protein Gel, 18 well, 30 μl | Bio Rad | Cat#: 3450124 |
| Amersham™ Protran® Western blotting membranes, nitrocellulose | Milipore Sigma | Cat#: GE10600002 |
| Corning ® 96-well clear flat bottom polystyrene non-treated microplate | Corning | Cat#: 3370 |
| Pierce™ Peptide Desalting Spin Columns | Thermo Scientific | Cat#: 89852 |
| Nalgene™ Oak Ridge High-Speed PPCO Centrifuge Tubes | Thermo Scientific | Cat#: 3119-0050PK |
| Falcon™ Round-Bottom Polysytrene Test Tubes with Cell Strainer Snap Cap, 5 mL | Thermo Scientific | Cat#: 08-771-23 |
